# Pathogenic role of cGAS-STING pathway in Duchenne muscular dystrophy

**DOI:** 10.64898/2026.11.13.766546

**Authors:** Alice Rannou, Nancy Manriquez, Sankalpa Chakraborty, Adarsh Balaji, Jonah Zarrow, Jocelyn Alfaro, Liang Li, Mario Fournier, Weixin Liu, Sophia Kangavari, Ahmed Ibrahim, Eduardo Marbán, Russell G. Rogers

**Author notes:** Address Correspondence to: Russell G. Rogers, PhD, Associate Professor, Smidt Heart Institute, Cedars-Sinai Medical Center, 8700 Beverly Boulevard, 1088 Davis Research Building, Los Angeles, California 90048.

## Abstract

The cGAS-STING pathway is a critical component of the innate immune response to infection and cell stress. Chronic activation underlies human diseases such as lupus and inflammatory bowel disease, but cGAS-STING has not been implicated in Duchenne muscular dystrophy (DMD). Compared to healthy controls, macrophages from DMD patients have high levels of cytosolic DNA and low levels of the 3’ DNA exonuclease, TREX1, a scavenger of cytosolic DNA upstream of cGAS-STING. Likewise, cytosolic DNA is elevated in macrophages from dystrophin-deficient *mdx* mice. Such macrophages over-produce 2’3’-cGAMP and secrete it locally, activating STING in neighboring myoblasts and thereby suppressing myogenesis. TY1, a synthetic non-coding RNA drug that enhances TREX1, attenuates cGAS-STING and restores cellular and tissue homeostasis in *mdx* mice. When administered intravenously or orally, TY1 not only halted disease progression but actually reversed cardiac and skeletal muscle pathology. Thus, pathological activation of cGAS-STING by cytosolic DNA mediates disease progression in DMD. Our work identifies TREX1, cGAS and STING as novel therapeutic targets in muscular dystrophy.

## INTRODUCTION

Duchenne muscular dystrophy (DMD), the most common heritable disease of muscle (1), arises from mutations in the *Dmd* gene, with eventual degeneration of both cardiac (2) and skeletal muscle. Loss of dystrophin, the *Dmd* gene product, increases cell membrane fragility and thereby fosters inflammation and fibrosis (3). In other settings associated with inflammation, such as infection or autoimmunity, DNA damage plays a key role: the release of fragmented double-stranded DNA (dsDNA) into the cytosol activates cyclic-GMP-AMP synthase (cGAS) (4), which, in turn, triggers the production of type I interferons and pro-inflammatory cytokines via cGAS-Stimulator of Interferon Genes (STING). However, DNA damage and cGAS-STING activation have not been implicated in DMD pathogenesis. Here, we report excess cytosolic dsDNA accumulation, and low levels of the 3’ DNA exonuclease TREX1, in *mdx* mouse and human DMD macrophages. Dystrophic macrophages are replete with 2’3’-cyclic-GMP-AMP (2’3’-cGAMP), a cyclic dinucleotide that not only activates STING endogenously but also traverses the cell membrane and functions as a paracrine activator of STING. Consumption of 2’3’-cGAMP by skeletal myoblasts markedly suppresses proliferation and fusion—key cellular processes required for muscle regeneration. Up-regulation of TREX1 by a synthetic non-coding RNA, TY1 (5), reduces cytosolic dsDNA accumulation, inhibits cGAS activation, and promotes an anti-inflammatory phenotype in *mdx* macrophages. Modulation of macrophage function by TY1 leads to systemic improvements in exercise performance, skeletal myopathy, and cardiomyopathy when administered intravenously or orally (6) to *mdx* mice. Collectively, our findings identify the cGAS-STING pathway as pathogenic in DMD, pinpointing novel opportunities for therapeutic intervention.

## RESULTS

### Cytosolic dsDNA accumulation in human DMD macrophages

Mounting evidence implicates cGAS-STING in the pathophysiology of diverse disease processes (7), including autoimmunity (8) and aging (9). Given that inflammation is a prominent in DMD (10), we wondered if cGAS-STING might play a role. We first sought to determine if human DMD macrophages, cultured from peripheral blood mononuclear cells in cytokine-containing media (11), accumulate cytosolic dsDNA. Relative to healthy age- and sex-matched donors, DMD macrophages contained more cytosolic dsDNA, evident in representative micrographs (Fig. 1A). Not only are DMD macrophages twice as likely to have cytosolic dsDNA, but also such cells contain more cytosolic dsDNA foci than healthy macrophages (Fig. 1B). Consistent with the increase in cytosolic dsDNA, TREX1, a 3’ DNA exonuclease responsible for degrading cytosolic dsDNA (12), is dramatically suppressed in DMD macrophages (Fig. 1C)—rationalizing further interrogation of cGAS-STING. To do so, we focused on *mdx* mice, where it was possible to avoid confounding effects of guideline-driven chronic corticosteroid use in DMD donors (13) (which can, in turn, suppress cGAS-STING (14, 15)).

**Figure 1:**
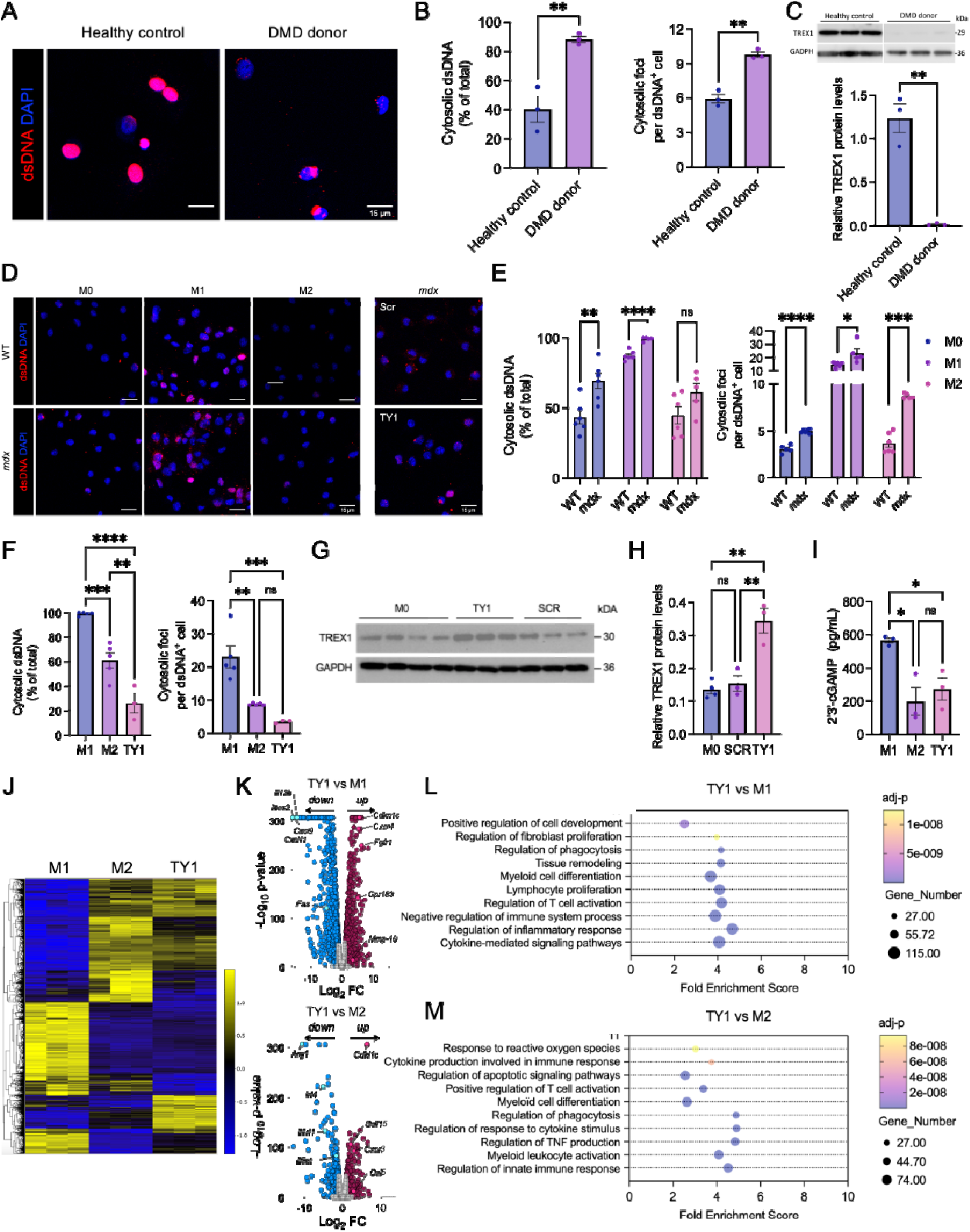
Cytosolic dsDNA accumulation activates cGAS in human DMD and *mdx* mouse macrophages. Human DMD and healthy control peripheral blood mononuclear cells were plated and cultured for 7 days in presence of human m-CSF (50 ng/mL). At day 7, M0 macrophages were obtained. The percentage of cells containing cytosolic dsDNA+ foci and the number of positive dsDNA+ foci per cell were quantified (**A&B**). Western blot of TREX1in M0 macrophages (**C**). Wild-type and *mdx* bone marrow derived-macrophages were isolated and cultured for 7 days in presence of m-CSF (10 ng/mL). At day 7, M0 macrophages were polarized to M1 (LPS 0.1 µg/ml; IFN-γ 50 ng/mL), M2 (IL13 10 ng/mL; IL4 10n g/mL), or conditioned by TY1 or scramble (50 nM) for 24 hours. The percentage of cells containing cytosolic dsDNA+ foci and the number of positive dsDNA+ foci per cell were quantified (**D, E&F**). Western blot of TREX1 in TY1 or scramble conditioned macrophages compared to M0 (inactivated) (**G&H**). 2’3’- cGAMP from the cell lysate of M0 and TY1-transfected macrophages using standard ELISA (**I**).RNA was collected and sequenced. Heatmap showing transcripts up- or down-regulated after exposure to TY1 and compared to M1 and M2 phenotypes (**J**). Volcano plot of the top differentially expressed genes in TY1-conditioned macrophages compared to M1(**K**; top) and M2 (**K**; bottom) phenotypes. Selected pathways enriched in TY1-conditioned macrophages compared to M1 (**L**) and M2 (**M**) phenotypes. Scale bars: 15 µm. N = 3-6 per group. Data are represented as mean ± SEM. Statistical significance was determined by ANOVA. *p < 0.05, **p < 0.01, ***p < 0.001, ****p < 0.0001, ns = not significant.

### Cytosolic dsDNA and cGAS-STING activation in *mdx* macrophages

We assessed the levels of cytosolic dsDNA in polarized *mdx* macrophages compared to naïve macrophages. When stimulated with pro-inflammatory cytokines (e.g., M1), mouse macrophages had elevated levels of cytosolic dsDNA compared to naive macrophages (e.g., M0) or macrophages exposed to anti-inflammatory cytokines (e.g., M2; Fig. 1D). These findings are not surprising: pro-inflammatory activation of macrophages induces accumulation of cytosolic dsDNA, perhaps driven by elevated production of reactive oxygen species (16). But the consistent increase in dsDNA in *mdx* macrophages, regardless of polarization state, is unexpected. Compared to macrophages from wild-type mice, similarly-activated *mdx* macrophages had greater levels of cytosolic dsDNA (Fig. 1D). Not only are *mdx* macrophages more frequently dsDNA+, but such dsDNA+ *mdx* macrophages also contain more dsDNA foci (Fig. 1E), verified to be cytosolic (i.e., not nuclear or mitochondrial) by DAPI and COX4 staining (SFig. 1A). Thus, for as-yet unexplored reasons, *mdx* macrophages appear to be pathologically susceptible to DNA damage (17), which may contribute to immune dysfunction—a hallmark of DMD (10).

To better understand the role of DNA damage in DMD, we tested the effects of a novel RNA drug (an exomer), TY1, that enhances the expression of TREX1 (5). Exposure to TY1 dramatically lowered the levels of cytosolic dsDNA in *mdx* macrophages, whether compared to pro-or anti-inflammatory polarization states (Fig. 1D). The percentage of cytosolic dsDNA+ *mdx* macrophages, and the number of dsDNA foci within such macrophages, were dramatically lower in *mdx* macrophages conditioned by TY1 (Fig. 1F). Unlike a scrambled control RNA oligonucleotide (SCR, with an identical nucleotide content but a sequence randomized to lack complementarity with mammalian genomes), TY1 boosts the protein levels of TREX1 in *mdx* macrophages (Fig. 1G&H). Because SCR had no effect on TREX1 protein levels, the bioactivity of TY1 is attributable to its unique sequence. To verify that TY1 lowers cytosolic dsDNA by upregulating TREX1, we transfected pro-inflammatory *mdx* macrophages (i.e., those with the greatest level of cytosolic dsDNA) with *Trex1* plasmid DNA and measured cytosolic dsDNA by immunocytochemistry. Western blot confirmed transfection with *Trex1* boosted protein levels of TREX1 (SFig. 1B). Relative to control, TREX1 up-regulation by *Trex1* lowered the percentage of dsDNA+ *mdx* macrophages and the number of dsDNA foci within such macrophages (SFig. 1C), reproducing what we observed with TY1 (Fig. 1F). As expected with excess cytosolic dsDNA, pro-inflammatory *mdx* macrophages produce high levels of 2’3’-cGAMP, a second messenger and activator of the pro-inflammatory STING pathway, which is 3-fold lower in anti-inflammatory (M2) *mdx* macrophages and those conditioned by TY1 (Fig. 1I). Cultured *mdx* macrophages take up TY1: a fluorescent form of TY1 (SFig. 1D) (18) accumulates in cells as seen either by flow cytometry (SFig. 1E) or confocal microscopy (SFig. 1F). Collectively, these findings demonstrate cGAS-STING is active in *mdx* macrophages and further motivate exploration of this pathway in DMD.

### Anti-inflammatory effects of TY1 in *mdx* macrophages

Chronic activation of cGAS-STING leads to dysregulated immune responses and propagation of pro-inflammatory signaling (19). In contrast, ablation of cGAS increases the abundance of wound-healing (anti-inflammatory) macrophages (20). Because TY1 lowers cGAS activity in *mdx* macrophages (Fig. 1I), we sought to determine if TY1 alters the expression of pro- and anti-inflammatory genes. Heatmaps of RNA-seq data show the transcriptome of TY1-conditioned *mdx* macrophages closely aligns with that of M2 anti-inflammatory macrophages (Fig. 1J), as verified by qPCR (SFig. 1G). Notably, TY1 boosted transcript levels of the anti-inflammatory cytokine, *Il10*, 5-fold relative to M2 anti-inflammatory macrophages (SFig. 1G), exemplifying broader differences in gene expression (Fig. 1K). Relative to M1 *mdx* macrophages, TY1-conditioned *mdx* macrophages express *Cxcr4* (21), *Gpr183* (22), *Cdkn1c* (23), and *Fgfr1* (24)—genes involved in macrophage chemotaxis and reparative function—and suppress *Cxcl9* (25), *Cxcl11* (26), *Fas* (27), *Il12b* (28), and *Nos2* (29)—genes involved in macrophage chemotaxis and inflammation. Relative to M2 *mdx* macrophages, TY1-conditioned *mdx* macrophages express *Cxcr3* (30), *Ccl5* (31)*, Cdkn1c* (23) and *Gdf15* (32)—genes involved in macrophage chemotaxis and tissue remodeling—and suppress the pro-fibrotic genes *Arg1* (33), *Il1rl1* (34), *Irf4* (35), and *Il6st* (36). Gene Ontology (GO) analysis of RNA-seq data revealed broad immune modulation of *mdx* macrophages by TY1 (Fig. 1 L&M). Thus, TY1 suppresses cGAS-STING and attenuates inflammatory gene signatures in *mdx* macrophages.

### TY1 improves function and structure in *mdx* mice

The immunomodulatory effects of TY1 on isolated macrophages motivated us to look for therapeutic benefits *in vivo*. Because cardiosphere-derived cells (37, 38) and their extracellular vesicles (37, 39–41) are effective when delivered intravenously, we infused TY1 in 9-12 month-old *mdx* mice, an age at which both skeletal myopathy and cardiomyopathy are evident (37, 41) (see Fig. 2A for experimental design). After 8 weeks, *mdx* mice that had received TY1: a) were able to run 20% farther than at baseline before exhaustion (Fig. 2B), b) produced 10% more isometric skeletal muscle torque than at baseline, whereas vehicle-infused controls were 50% weaker (Fig. 2C), and c) showed preserved cardiac function, whereas controls decreased by 20% (Fig. 2D). The decreases in skeletal muscle and heart function in *mdx* mice infused with vehicle over the 8 weeks of observation reflect the natural history of late-stage DMD (37, 41). Underlying the functional improvements, histology revealed striking reductions of dystrophic features (Fig. 2E). Relative to control, skeletal muscle and hearts of *mdx* mice that had received TY1 showed decreased interstitial fibrosis (Fig. 2E&F). Skeletal muscle from *mdx* mice that had received TY1 also exhibited twice as many myofibers per cross-sectional area (Fig. 2G), hinting TY1 may promote myogenesis—a possibility we explore later. Similar benefits were seen when we fed *mdx* mice TY1 in a casein-chitosan oral formulation (6) (see Fig. 2H for experimental design): exercise performance improved (Fig. 2I), skeletal muscle function was preserved (Fig. 2J), and heart function was augmented (Fig. 2K), in contrast to controls. Histology revealed similar improvements in fibrosis, and increases in myofiber numbers, with oral TY1 as compared to intravenous delivery (Fig. 2L-M). Collectively, these findings indicate TY1, whether delivered intravenously or orally, exerts striking disease-modifying bioactivity in aged *mdx* mice.

**Figure 2:**
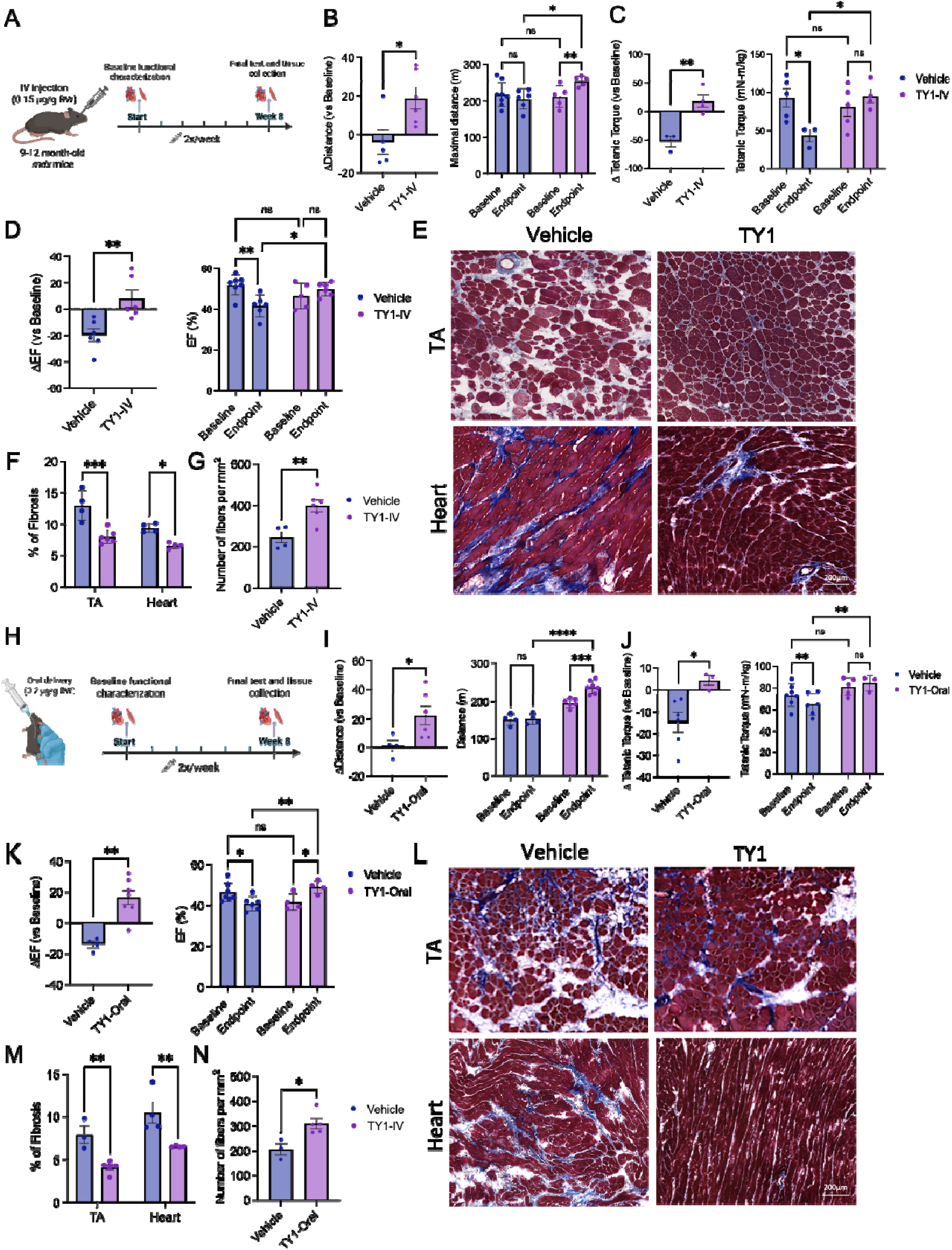
Functional and structural benefits of TY1 in *mdx* mice. Schematic of experimental design delivering TY1 by intravenous infusion (**A**). Mice received vehicle or TY1 (0.15 µg/kg body weight) twice-weekly via retro-orbital injection for 8 weeks. Functional measurements were made at baseline and endpoint. Histological assessment was made at endpoint. Maximal exercise capacity (**B**), isometric tetanic torque (**C**), and ejection fraction (**D**) were improved in *mdx* mice that had received TY1. Analysis of Masson’s trichrome staining showed TY1 lowered fibrosis in the skeletal muscle and heart, and increased myofiber numbers in the tibialis anterior (TA) (**E-G**). Schematic of experimental design delivering TY1 by oral gavage (**H**). Mice received vehicle or TY1 (0.2 µg/kg body weight) twice-weekly via oral gavage for 8 weeks. Functional measurements were made at baseline and endpoint. Histological assessment was made at endpoint. Maximal exercise capacity (**I**), isometric tetanic torque (**J**), and ejection fraction (**K**) were improved in *mdx* mice that had received TY1. Analysis of Masson’s trichrome staining showed TY1 lowered fibrosis in the skeletal muscle and heart, and increased myofiber numbers in the TA (**L-N**). Scale bars: 200 µm. N = 4-6 per group. Data are represented as mean ± SEM. Statistical significance was determined by ANOVA or t-test. *p < 0.05, **p < 0.01, ***p < 0.001, ****p < 0.0001, ns = not significant.

### cGAS-STING is active in *mdx* skeletal muscle

Although DNA damage has been observed in dystrophic skeletal muscle (42), the cGAS-STING pathway has yet to be probed. Relative to wild-type skeletal muscle, cGAS, phospho-TBK1, total TBK1, and phospho-IKKα/β were elevated at the protein level in *mdx* skeletal muscle (Fig. 3A&B). In addition, 2’3’-cGAMP was more plentiful in *mdx* than in wild-type skeletal muscle (Fig. 3C). All these indicators of cGAS-STING activity were suppressed by oral TY1 (Fig. 3A-C). Meanwhile, TY1 decreased the proportion of pro-inflammatory (CD80+), and increased the proportion of anti-inflammatory (CD206+), macrophages while not affecting total macrophage number (i.e., CD68+; Fig. 3D&E). Consistent with our earlier findings *in vitro* (Fig. 1), cGAS-STING suppression *in vivo* shifts macrophages towards an anti-inflammatory phenotype that is known to support tissue repair (19).

**Figure 3:**
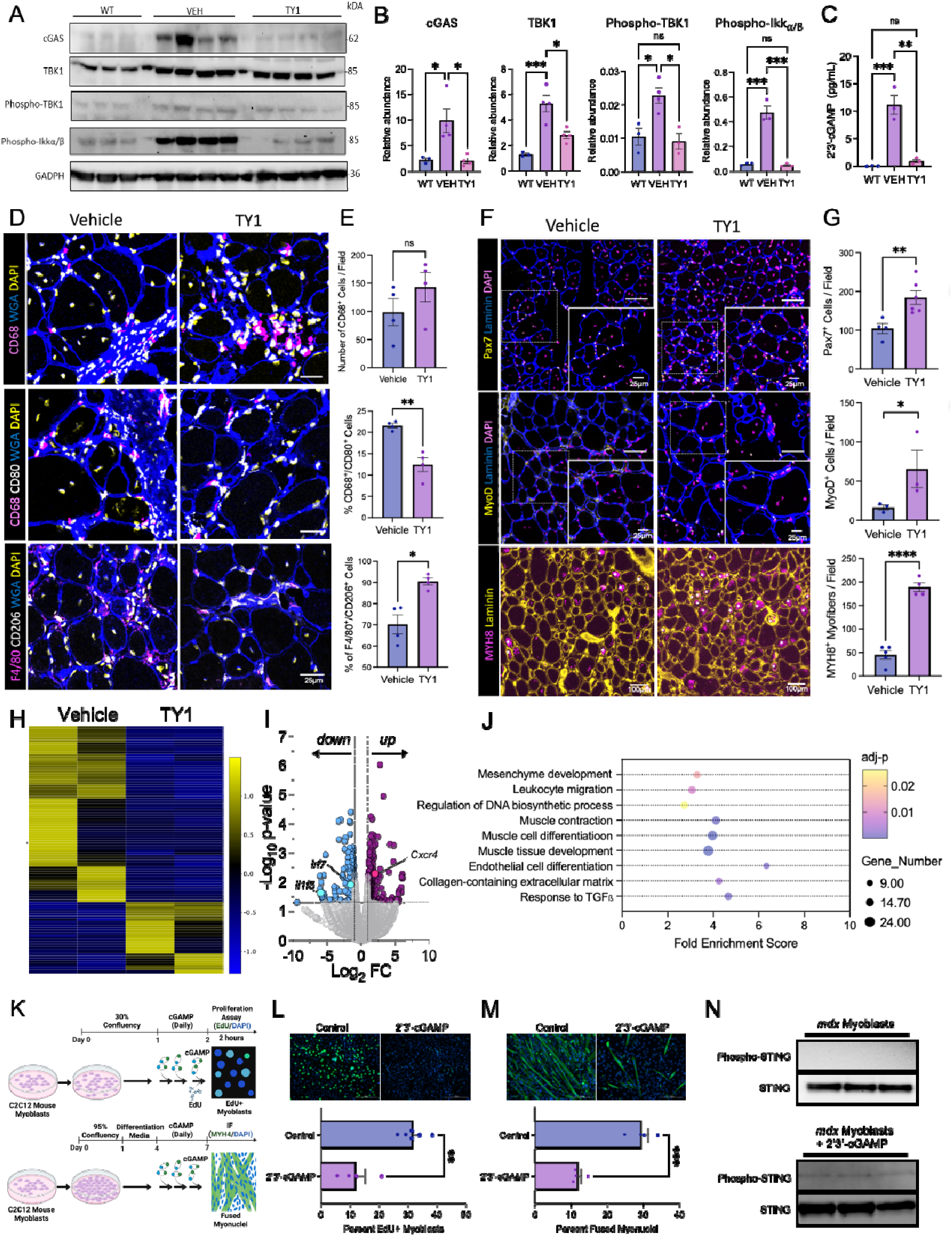
cGAS-STING suppresses skeletal myogenesis. Western blot of tibialis anterior (TA) lysates show cGAS, TBK1, phospho-TBK1, and phospho-Ikkα/β are increased in *mdx* mice but lowered to wild-type levels by TY1 (**A&B**). ELISA of TA lysates show 2’3’-cGAMP levels are increased in *mdx* mice but lowered to wild-type levels by TY1 (**C**). Immunohistochemistry of TA sections show TY1 had no effect on the global macrophage population (CD68+, top), but decreased pro-inflammatory (CD68+/CD80+, middle) and increased anti-inflammatory (CD68+/CD206+, bottom) macrophages (**D&E**). Immunohistochemistry of TA sections show TY1 elevated the number of Pax7+ cells (top), MyoD+ cells (middle), and neonatal myosin heavy chain (MYH8)+ myofibers (bottom) (**F&G**). Bulk RNA-seq show transcriptome-wide changes gene expression changes in *mdx* skeletal muscle (**H**). Volcano plot shows the top up- and down-regulated genes by TY1 (**I**). GO indicated robust enrichment of pathways related to skeletal muscle myogenesis (**J**). Schematic of experimental design testing the effects of 2’3’-cGAMP on C2C12 myoblast proliferation and myotube formation (**K**). Immunocytochemistry of proliferating (EdU+) C2C12 myoblasts show 100 μM 2’3’- cGAMP suppressed nuclear incorporation of EdU (**L**). Immunocytochemistry of differentiating (MHY4+) C2C12 myotubes show 100 μM 2’3’-cGAMP blocked myonuclei fusion into myotubes (**M**). Western blot of *mdx* mouse myoblasts shows 100 μM 2’3’- cGAMP stimulates STING phosphorylation (**N**). Scale bars: 200 μm, 100 μm, 50 μm, or 25 μm. N = 2-6 per group. Data are represented as mean ± SEM. Statistical significance was determined by ANOVA or t-test. *p < 0.05, **p < 0.01, ****p < 0.0001, ns = not significant. Asterix in **D** (bottom) denote MYH8+ myofibers.

### cGAS-STING crosstalk suppresses skeletal muscle myogenesis

Macrophages are key regulators of myogenesis, by crosstalk with muscle stem cells (43). To test if chronic elevations in cGAS-STING might impair myogenesis, we quantified Pax7+ muscle precursor cells, MyoD+ myoblasts, and MYH8+ regenerating myofibers with or without TY1. Relative to control (i.e., vehicle-exposed *mdx* skeletal muscle with elevated cGAS-STING), TY1 boosted the numbers of Pax7+ and MyoD+ myogenic cells, and MYH8+ regenerating myofibers (Fig. 3F&G), supporting the idea that cGAS-STING suppresses myogenesis. Heatmaps of bulk RNA-seq of *mdx* skeletal muscle show major transcriptomic changes induced by TY1, reflecting the complexity of cGAS-STING (44), the immune microenvironment, and myogenesis (Fig. 3H). In particular, TY1 up-regulates *Cxcr4*, a gene involved in satellite cell activation, expansion, and self-renewal (45), and down-regulates the expression of *Irf7* (Fig. 3I), a downstream target of the cGAS-STING pathway (46), and *Il1f5*, a pro-inflammatory cytokine involved in fibrosis (47). GO analysis further revealed enrichment of various pathways involved in myogenesis (e.g., “muscle tissue development” and “muscle cell differentiation”; Fig. 3J) with TY1.

These data motivated us to further examine cGAS-STING crosstalk between *mdx* macrophages and myoblasts. We first verified *mdx* macrophages secrete 2’3’- cGAMP, which is lowered by TY1 (SFig. 3A). We then exposed C2C12 myoblasts to varying concentrations of 2’3’-cGAMP to confirm uptake and STING activation. Indeed, STING phosphorylation increased as a function of 2’3’-cGAMP concentration (with 100 µM stimulating the greatest phosphorylation); this effect was observed whether or not 2’3’-cGAMP was packaged in a transfection reagent (SFig. 3B; cf. SFig. 3C). We next probed the effect of STING activation on C2C12 myoblast proliferation and myotube formation (see Fig. 3K for experimental design). EdU pulse-chase revealed 100 µM 2’3’- cGAMP decreased C2C12 myoblast proliferation by 62 % (31% vs 12%; Fig. 3L). Further, myosin heavy chain 4 (MYH4) immunocytochemistry showed 100 µm 2’3’- cGAMP decreased myonuclei fusion into myotubes by 59% (29% vs 12%; Fig. 3M). In primary *mdx* myoblasts, 2’3’-cGAMP, but not control, induced STING phosphorylation (Fig. 3N). Taken together, these data demonstrate cGAS-STING crosstalk dampens the myogenic actions of skeletal myoblasts, likely contributing to the regenerative defects characteristic of DMD.

### Reparative remodeling is blocked by cGAS-STING in *mdx* hearts

Inflammatory/fibrotic cardiomyopathy is a deadly manifestation of advanced DMD (2). We therefore wondered if elevated cGAS-STING underlies inflammatory pathology in hearts, as it does in skeletal muscle. Relative to wild-type hearts, cGAS and phospho-IKKα/β protein levels were modestly elevated in *mdx* hearts, but decreased by TY1 (Fig. 4A&B). 2’3’-cGAMP levels were also modestly increased in *mdx* hearts and lowered by TY1 (Fig. 4C). It is currently unclear why cGAS-STING activity in the *mdx* mouse heart is less robust than in skeletal muscle, but dissection of the differences is beyond the scope of this report. In any case, as with DMD macrophages, TREX1 is lower in *mdx* hearts but is increased to wild-type levels by TY1 (Fig. 4B). Together, these data suggest cGAS-STING may be active in *mdx* hearts and can be targeted by TY1. Unlike *mdx* skeletal muscle, TY1 decreased the number of CD68+ macrophages (Fig. 4D&E), which may reflect an underlying dichotomy between the two tissues (48). Similar to the effects in *mdx* skeletal muscle, however, TY1 decreased the proportion of CD80+ (pro-inflammatory), and increased the proportion of CD206+ (anti-inflammatory), macrophages (Fig. 4D&E), reflecting comparable immunomodulatory actions of cGAS-STING suppression. Heatmaps of bulk RNA-seq data from *mdx* hearts show major transcriptomic changes induced by TY1, consistent with our observations in *mdx* skeletal muscle (Fig. 4F). TY1 down-regulates *Meg3* and *Cd226* and up-regulates *Nrg1*, *Myh4*, and *Myh1* (Fig. 4G); silencing of *Meg3* and *Cd226* have been shown to block cardiomyocyte apoptosis (49) and skew macrophage polarization towards an anti-inflammatory phenotype (50), respectively, while *Nrg1* up-regulation promotes cardiomyocyte proliferation (51). Changes in the expression of *Meg3*, *Cd226*, and *Nrg1* may contribute to the TY1-induced structural improvements in *mdx* hearts (Fig. 2E&L). GO analysis revealed high enrichment scores for “cardiomyocyte differentiation”, “muscle tissue development”, “regulation of angiogenesis”, and “response to TGF-β” (Fig. 4H). Immunohistochemistry verified key predictions of the GO analysis: hearts from TY1-exposed *mdx* mice showed more actively cycling/dividing (i.e., Ki67+) cardiomyocytes (Fig. 4I&J), indicating increased cell cycle activity relative to control (52); moreover, CD31 was 33% greater in area after TY1 (Fig. 4K&L), as expected by increased angiogenesis. Collectively, these data support the concept that cGAS-STING activity inhibits tissue repair and homeostasis in DMD heart; conversely, suppression of cGAS-STING by TY1 liberates the reparative response and improves cardiac function (cf. Fig. 2).

**Figure 4:**
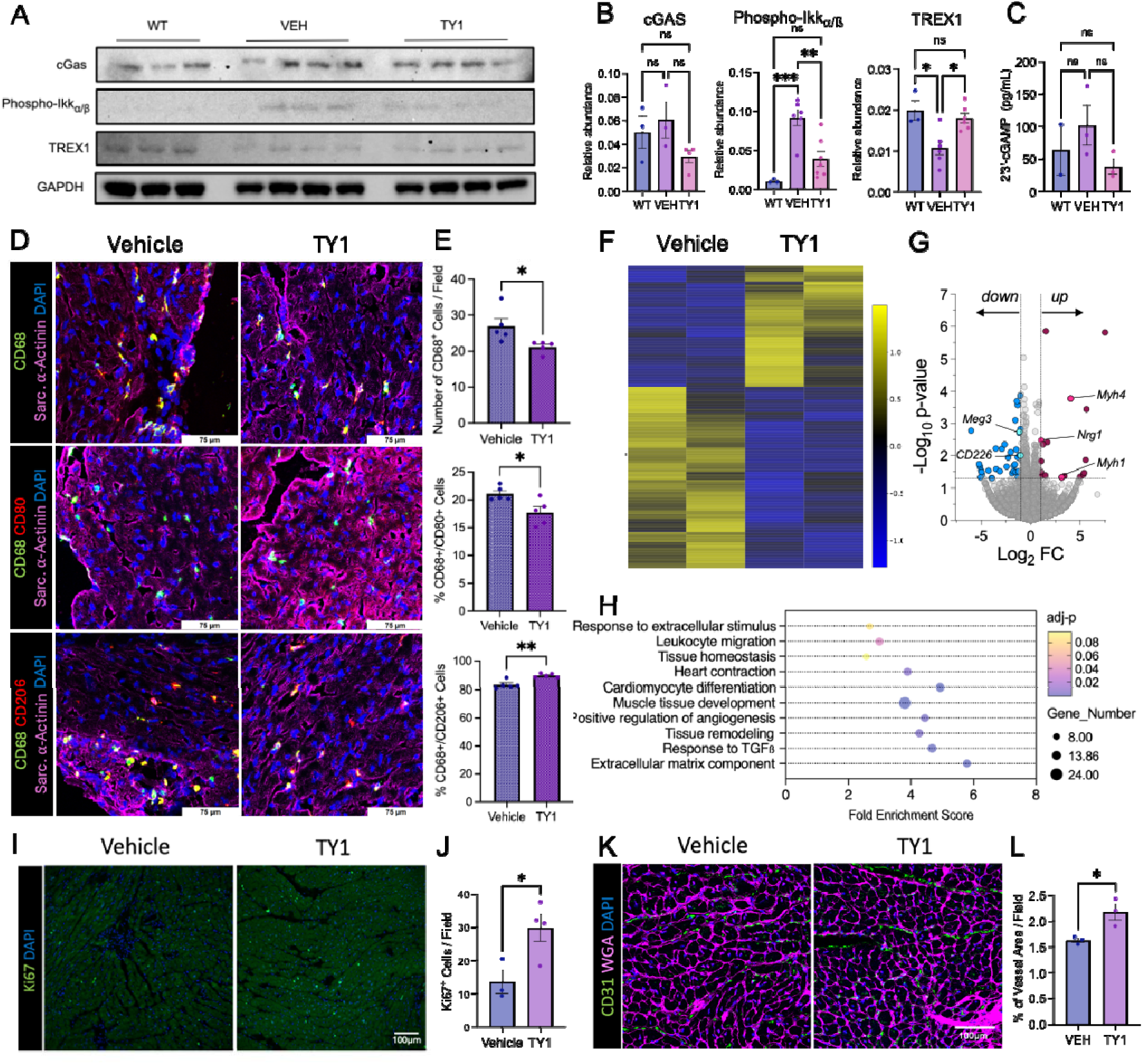
cGAS-STING regulates reparative remodeling in *mdx* hearts. Western blot of heart lysates show TREX1 is lowered in *mdx* mice but increased to wild-type levels by TY1, while cGAS and phospho-Ikkα/β is increased in *mdx* mice but lowered to wild-type levels by TY1 (**A&B**). ELISA of heart lysates show 2’3’-cGAMP levels tend to be higher in *mdx* mice but lowered by TY1 (**C**). Immunohistochemistry of heart sections show TY1 lowered global macrophage accumulation (CD68+, top) and pro-inflammatory (CD68+/CD80+, middle) macrophages, and increased anti-inflammatory (CD68+/CD206+, bottom) macrophages (**D&E**). Bulk RNA-seq show transcriptome-wide changes gene expression changes in *mdx* hearts (**F**). Volcano plot shows the top up- and down-regulated genes by TY1 (**G**). GO indicated robust enrichment of pathways related to tissue remodeling and angiogenesis (**H**). Immunohistochemistry of heart sections show TY1 elevated the number of cycling Ki67+ cells (**I&J**) and CD31+ vessel area (**K&L**). Scale bars: 100 μm. N = 3-5 per group. Data are represented as mean ± SEM. Statistical significance was determined by ANOVA or t-test. *p < 0.05, **p < 0.01, ns = not significant.

### Effects of cGAS-STING suppression by TY1 are macrophage-dependent

The data, thus far, suggest that macrophages figure prominently in the disease-modifying bioactivity of TY1 (consistent with evidence that inflammatory macrophages underlie inflammation-related pathology in DMD (53)), but fall short of establishing a cause-effect relationship. To determine if macrophages are the primary cellular target of TY1, we created an inducible macrophage-depletable *mdx* mouse model by crossing *mdx* and MaFIA mice, which harbor a suicide gene under control of the *c-fms* promoter—enabling effective depletion of circulating and tissue macrophages (54) (SFig. 4A&B). No differences were noted in heart function among wild-type, *mdx* or *mdx*/MaFIA mice at ∼2 months of age (SFig. 4C); while heart function and exercise performance declined between 2 and 8 months of age in *mdx*/MaFIA mice (SFig. 4D&E), the changes are similar to those in non-MaFIA *mdx* mice (41). In terms of macrophage phagocytic activity, *mdx* and *mdx*/MaFIA mice were comparable (SFig. 4F). Nevertheless, exposure of *mdx*/MaFIA mice to AP20187 triggered high mortality (SFig. 4G), hinting at a central homeostatic role for macrophages.

Our experimental protocol (Fig. 5A) compared responses to TY1 in macrophage-intact versus macrophage-depleted *mdx*/MaFIA mice (the latter being limited to hardy survivors of the depletion process). In immunocompetent mice that received TY1, muscle torque was 20% greater than at baseline, while torque in controls decreased by 20% (Fig. 5B). Meanwhile, global heart function went up 12%, whereas controls decreased by 8% (Fig. 5C). In contrast to macrophage-competent mice, TY1 had no major effect on muscle (Fig. 5B) or heart (Fig. 5C) function in macrophage-depleted *mdx*/MaFIA mice. Histology revealed an unexpected finding—skeletal muscle (Fig. 5D&E) and hearts (Fig. 5F&G) were more fibrotic when macrophages were not present, regardless of intervention; increased cardiac fibrosis predisposes to sudden death (55), perhaps explaining the high mortality associated with macrophage depletion. However, when macrophages were intact, TY1 was effective at reducing interstitial fibrosis in skeletal muscle and heart (Fig. 5D-G), in line with our findings in non-MaFIA *mdx* mice (cf. Fig. 2). To further verify TY1’s disease-modifying bioactivity is macrophage-dependent, we tested the efficacy of TY1 in *mdx* mice with or without macrophage depletion by clodronate liposomes(56) (see SFig. 5A for experimental design). Consistent with data acquired in *mdx*/MaFIA mice, skeletal muscle (SFig. 5B) and heart (SFig. 5C) function were lower in *mdx* mice that had received TY1 without macrophages compared to *mdx* mice with macrophages. Further, interstitial fibrosis of skeletal muscle (SFig. 5D&E) and heart (SFig. 5F&G) were higher in *mdx* mice that had received TY1 without macrophages, compared to *mdx* mice with macrophages.

**Figure 5:**
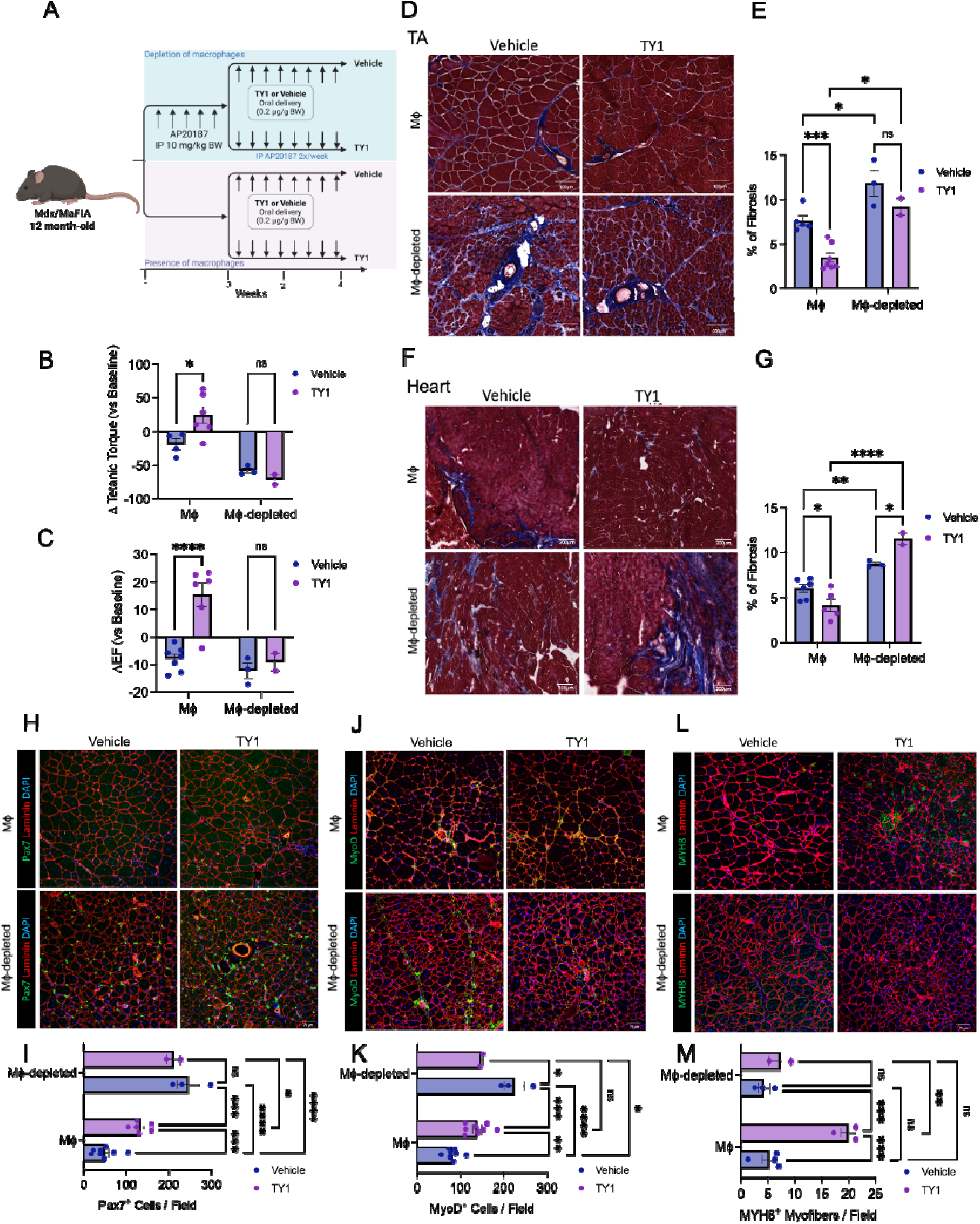
Macrophage-dependent effects of cGAS-STING suppression. Schematic of experimental design to deplete circulating and tissue macrophages (**A**). MaFIA mice harboring the *mdx* mutation were administered AP20187 (10 mg/kg, intraperitoneal) for 5 consecutive days. Macrophage depletion was maintained by twice-weekly injections of AP20187. Following macrophage depletion, mice received vehicle or TY1 (0.2 µg/kg body weight) twice-weekly by oral gavage for 4 weeks. Functional measurements were made at baseline (after depletion but before intervention) and endpoint. Histological assessment was made at endpoint. Isometric tetanic torque (**B**) and ejection fraction (**C**) were improved by TY1 but only when macrophages were present. Analysis of Masson’s trichrome staining showed fibrosis was only lowered by TY1 in the tibialis anterior (TA) (**D&E**) and heart (**E&F**) when macrophages were present. Immunohistochemistry of TA sections show TY1 elevated the number of Pax7+ cells (**H&I**), MyoD+ cells (**J&K**), and MYH8+ (**L&M**), but only when macrophages were present. Scale bars: 200 μm or 75 μm. N = 2-6 per group. Data are represented as mean ± SEM. Statistical significance was determined by ANOVA. *p < 0.05, **p < 0.01, ***p < 0.001, ****p < 0.0001, ns = not significant.

Analysis of myogenic cells in skeletal muscle of *mdx*/MaFIA mice revealed major differences with macrophage depletion. When macrophages were intact, TY1 increased the number of Pax7+ muscle precursor cells (Fig. 5H&I), MyoD+ myoblasts (Fig. 5J&K), and MYH8+ myofibers (Fig. 5L&M). In contrast, when macrophages are depleted, Pax7+ (Fig. 5H&I) and MyoD+ (Fig. 5J&K) cells accumulate but fail to form MYH8+ regenerating myofibers (Fig. 5L&M). Thus, the effect of TY1 is dependent on macrophages, but global macrophage depletion leads to a dramatic increase in the number of myofibers (SFig. 6A). Size distribution analysis showed no major effect of TY1 on myofiber size, but macrophage depletion shifted the histogram towards diminutive myofibers (e.g., 200-800 µm^2^) (SFig. 6B)—as if myogenic cells stall at the fusion/maturation stage in the absence of macrophages. Taken together, these findings provide compelling evidence that macrophages, whether circulating or tissue-resident, are the primary cellular targets of TY1. Such cells also prominently modulate myogenesis.

## DISCUSSION

DMD is a myodegenerative disease caused by underlying genetic mutations in the *Dmd* gene, leading to deficiency of dystrophin —the major cytoskeletal protein supporting sarcolemmal stability. Although the root cause of DMD is dystrophin deficiency, tissue damage resulting from contraction of dystrophin-deficient muscle induces inflammation. Chronic and dysregulated inflammation is a known driver of DMD disease progression, but the molecular mechanism is less clear. Here, we have shown the pro-inflammatory cGAS-STING pathway is active in macrophages, skeletal muscle, and hearts of *mdx* mice. This evolutionarily ancient innate immunity pathway has been well studied in other settings such as autoimmunity (8) and aging (9), but had not been recognized to play a role in DMD.

The non-coding RNA drug, TY1, counters cGAS-STING activity *in vitro* and *in vivo*, and reverses key phenotypic abnormalities of DMD. TY1—the prototype for a new class of RNA drugs known as exomers—works by enhancing TREX1, the gatekeeper to cGAS-STING activation. TREX1 degrades cytosolic dsDNA that may accumulate during DNA replication (57) or damage (58). Here, we show dsDNA accumulates in the cytosol of human DMD and *mdx* mouse macrophages, and, in turn, activates cGAS. Activation of cGAS generates 2’3’-cGAMP which turns on STING within the macrophage (up-regulating pro-inflammatory genes), or it can be secreted and taken up by neighboring cells. In myoblasts, activation of STING suppresses myogenesis, underlying the broad regenerative defects observed in DMD skeletal muscle. The previously unrecognized role of cGAS-STING in DMD now opens new opportunities for therapeutic intervention.

Not only does TY1 serve as a novel reagent to probe the effects of cGAS-STING activity, but the exomer may also be a promising therapeutic candidate to target dysregulated inflammation in DMD. Indeed, TY1 is the first RNA drug with oral bioavailability (6), a pragmatic advantage in the treatment of DMD (which typically is diagnosed in young needle-averse children). Although not intended to correct the underlying genetic defect of DMD, TY1 effectively targets the major cause of disease progression: inflammation. In so doing, TY1 reverses key disease manifestations of DMD, at least partly by potent myogenic effects on dystrophic skeletal muscle. In principle, an oral agent that works by attacking inflammation and fibrosis would not only be complementary to, but also potentially synergistic with, emerging gene therapies.

## METHODS

### Animals and Experimental Design

This study used homozygous female *mdx* (C57BL/10ScSn-*Dmd^mdx^*/J; Jackson Labs #001801) or wild-type (C57BL/10J; Jackson Labs #000665) mice between 9-12 months of age and a novel macrophage-depletable *mdx* mouse model—by crossing *mdx* and MaFIA (54) (Macrophage Fas-Induced Apoptosis, Jackson Labs #005070) mice. A colony of macrophage-depletable *mdx* mice was established and is maintained in the vivarium at Cedars-Sinai. Here, we do not explore the influence of sex, as DMD is X-linked and affects only males. Thus, we used the most consistent mouse model, female *mdx* mice bred to homozygosity (due to the more severe cardiac phenotype than male counterparts (59)); notably, female *mdx* mice have sufficed as the sole preclinical model to support three prior INDs (HOPE-Duchenne, HOPE-2, and HOPE-3). Mice were kept in a controlled environment under pathogen-free conditions with a 12-hour light-dark cycle and provided food and water *ad libitum*. All animal procedures were approved by the Institutional Animal Care and Use Committee at Cedars-Sinai. TY1 (0.15 µg/g body weight [intravenous] or 0.2 µg/g body weight [oral]) and controls were administered to mice by retro-orbital (i.e., intravenous) injection or oral gavage twice-weekly for 4-8 weeks. A 25% increase in dosage was added for oral delivery to offset decreased loading efficiency(6).

### Macrophage Depletion

*mdx*/MaFIA mice harbor a macrophage-driven *c-fms* promoter expressing eGFP and an inducible Fas-mediated suicide gene, which effectively depletes circulating and tissue resident cells of the monocyte/macrophage lineage. Macrophages were depleted by intraperitoneal injections of AP20187 (Takara Bio) for 5 consecutive days at a dose of 10 mg/kg. Macrophage depletion was maintained by twice-weekly injections of AP20187 (1 mg/kg) after the initial loading phase. Lyophilized AP20187 was reconstituted in 100% ethanol to a stock concentration of 62.5 mg/ml and stored at – 20°C. Injectable AP20187 was prepared fresh by diluting the AP20187 stock to 2.5 mg/ml in a solution of 4% ethanol, 10% PEG-400, and 1.7% Tween-20 in water. Volume was adjusted for mouse body weight to normalize the effective dose and averaged 130 µl per mouse. Macrophage depletion was confirmed in various tissues by flow cytometry. To pharmacologically deplete macrophages from *mdx* mice, Clodrosome (Encapsula NanoSciences) was diluted with sterile PBS (2:1) and 200 µL was injected intravenously via tail vein for 3 consecutive days. Macrophage depletion was maintained by twice-weekly injections of diluted Clodrosome.

### TY1 Synthesis and Formulation

TY1, a 24-nucleotide chemically-modified non-coding RNA oligonucleotide (60), was synthesized by Integrated DNA Technologies. Purity was assessed by mass spectrometry, and the sequence was verified. TY1 was complexed with DharmaFECT transfection reagent per the manufacturer’s recommended protocol (Perkin Elmer); this formulation was used for all *in vitro* and intravenous applications. To produce the oral formulation (6), a 5% bovine casein solution (in Ultrapure water) was added to the TY1-DharmaFECT complex at a ratio of 1:10 (v/v) and incubated at room temperature for 15 min. An equal volume of 0.1% acetic acid and 0.2% chitosan solution was then added to the mixture to form casein-chitosan micelles. The TY1:DharmaFECT:casein-chitosan micelle solution was further incubated at room temperature for 1 hour; this formulation was used for oral administration by gavage.

### Echocardiography

Following light isoflurane anesthesia, parasternal long-axis B-mode videos were captured using a Vevo 3100 system (Visual Sonics). Left ventricular ejection fraction was quantified using Visual Sonics v2.0.0 software. Measurements were taken from three or more consecutive cardiac cycles per mouse per time point.

### Graded Treadmill Exercise Test

Mice underwent a graded exercise test using an Exer-3/6 rodent treadmill (Columbus Instruments) at a 5° incline. During the acclimatization period, mice were left undisturbed for 30 min before the belt was engaged. Once the belt was activated, mice were encouraged (by gentle nudging) to familiarize themselves with walking on the treadmill at a pace of 10 m/min for an additional 20 minutes. Following acclimatization, the exercise protocol engaged, activating the shock grid at 0.15 mA with a frequency of 1 shock per sec. The belt was accelerated at 1 m/min per min until volitional exhaustion. Mice that rested on the shock grid for more than 10 seconds with gentle nudging were considered to have reached their maximal exercise capacity, and their accumulated distance traveled was recorded.

### *In Vivo* Skeletal Muscle Physiology

The torque-frequency relationship of the left anterior crural muscles (e.g., tibialis anterior, extensor digitorum longus, and extensor hallucis longus) was measured as previously described (61–64). Briefly, anesthesia was induced with 4% vaporized isoflurane and maintained with 1.5% vaporized isoflurane at a flow of 400 mL O_2_/min. Core temperature was maintained at 37°C using a temperature-controlled platform. The left knee was stabilized and the left foot was secured to an aluminum shoe connected to the shaft of an Aurora Scientific 300C servomotor and force transducer. Sterilized platinum needle electrodes (Natus Medical) were inserted percutaneously to stimulate the left common peroneal nerve. Stimulation voltage and electrode placement were optimized with 5–15 isometric contractions (200 ms train of 0.1 ms pulses delivered at 300 Hz). Isometric torque was recorded as a function of stimulation frequency (1–300 Hz).

### Mouse and Human Macrophage Culture and Transfection

Mouse bone marrow-derived monocytes/macrophages were derived from hematopoietic stem/progenitor cells, as described (65). Briefly, bone marrow was harvested by flushing mouse femurs with PBS (supplemented with 1% FBS, 2 mM EDTA) and passed through a 70-µm cell strainer. Red blood cells were lysed with ACK buffer (Invitrogen) and remaining bone marrow cells were washed with Iscove’s-Modified Dulbecco Media (IMDM, Gibco) supplemented with 10 ng/ml mouse M-CSF (R&D Systems). Macrophages were generated by resuspending bone marrow cells in IMDM supplemented with 10% FBS, 1% penicillin/streptomycin, and 10 ng/ml mouse M-CSF for 7 days. Cells were left untouched for 3 days, and then the culture media was changed every 2 days until day 7. Macrophages were then polarized to a pro-inflammatory (M1) phenotype (100 ng/ml LPS and 50 ng/ml IFN-γ; Sigma-Aldrich and R&D Systems, respectively), or to an anti-inflammatory (M2) phenotype (10 ng/ml IL-4 and IL-13; R&D Systems) for 24 hours. TY1 or scramble RNA control were prepared as described above, and 50 nM RNA was added to the macrophage culture medium for 24 hours. To overexpress TREX1 in *mdx* mouse macrophages, mouse *Trex1* plasmid DNA (MC209585, OriGene) or control (PS1000001, OriGene) was complexed with FuGENE HD transfection reagent (Promega), according to the manufacturer’s recommend protocol. *Mdx* macrophages were polarized with pro-inflammatory cytokines (as described above) and then incubated with *Trex1* plasmid DNA or control for 72 hours. Human peripheral blood mononuclear cells (PBMCs) were acquired from CureDuchenne Link (DMD donor) or purchased from Lonza (CC-2702, healthy control). DMD donors were on corticosteroid therapy at the time of collection. PBMCs were suspended in Mononuclear Cell Medium (PromoCell) for one hour at 37°C, and then plated in RPMI-1640 supplemented with 10% FBS, 1% penicillin/streptomycin, and 50 ng/ml human M-CSF (Invitrogen) for 7 days. PBMCs were left untouched for 3 days, and then the culture media was changed every 2 days.

### Mouse Myoblast Isolation and Culture

C2C12 myoblasts (ATCC) were cultured on 0.5% gelatin coated plates in Dulbecco’s-Modified Eagle Medium (DMEM, Gibco) supplemented with 20% FBS and 1% penicillin/streptomycin until reaching the appropriate confluency for assays. Primary mouse myoblasts were isolated from 4–6-week-old C57BL/10J (Jackson Labs #000665) or *mdx* mice. Briefly, hind limb muscles were dissected, minced, and digested with 0.2% collagenase II for 50 min at 37°C. Digesting tissues were intermittently agitated. Following digestion, tissues were triturated and passed through a 70-µM cell strainer. Isolated cells were pre-plated 3-5 times for 25 min to remove fibroblasts. Primary myoblasts were then plated on 10% Matrigel-coated plates and grown until experimental use.

### RNA Purification, Sequencing, and Quantitative Polymerase Chain Reaction

Isolated cells and tissues were washed with PBS and prepared for RNA isolation using the miRNeasy Mini Kit (Qiagen), according to the manufacturer’s recommended protocol. RNA concentration and purity were measured using a NanoDrop spectrophotometer (Thermo Fisher). RNA sequencing (RNA-seq) was performed as described(37). Briefly, a library was constructed using the TruSeq Stranded mRNA library preparation kit. Complementary DNA (cDNA) was synthesized from fragmented RNA and enriched by polymerase chain reaction (PCR). Sample libraries were multiplexed and sequenced on a NextSeq 500 platform (Illumina) using 75 base pair single-end sequencing. Raw reads were aligned to the mouse transcriptome using STAR/RSEM. Quantitative PCR (qPCR) was performed as described^16^. Briefly, cDNA was synthesized from 2 µg total RNA using a RevertAid first-strand cDNA synthesis kit (Thermo Fisher) and amplified using SsoAdvanced Universal SYBR Green Supermix (Bio-Rad) on a CFX Opus Real-Time PCR System (Bio-Rad). qPCR primers were designed using Primer-BLAST and are listed in Table S1. Raw RNA-seq data has been deposited in Gene Expression Omnibus: GSE282744 (*in vitro*) and GSE283913 (*in vivo*).

### Histology

At study endpoints, dissected tissues (e.g., tibialis anterior [TA] and heart) were embedded in Optimal Cutting Temperature (OCT, Sakura) compound and frozen in 2-methylbutane pre-cooled in liquid nitrogen (66). Serial cryosections (7-10 µm-thick) were cut using a Leica cryostat (CM3050S) and were adhered to Superfrost™ Plus microscope slides (Fisher Scientific). Cryosections were then post-fixed with 10% neutral-buffered formalin for 30 min at room temperature and processed for Masson’s trichrome staining using Weigert′s Iron Hematoxylin Set (HT1079, Sigma-Aldrich) and Trichrome Stain (Masson) Kit (HT15, Sigma-Aldrich), according to the manufacturer’s protocol. Histological slides were imaged using an Aperio AT Turbo slide scanner (Leica) at 40X magnification. Images were analyzed using ImageJ software (National Institutes of Health). Tissue fibrosis was quantified by dividing the area of blue staining (collagen) by the area of red staining (cytoplasm) of the entire tissue. ImageJ was also used to count myofiber numbers and to quantify the cross-sectional area (1000 myofibers per specimen).

### Immunohistochemistry

Specimens on cryosections were post-fixed with ice-cold acetone for 30 min at −20°C and then incubated with protein blocking buffer (Dako) for 1 hour at room temperature. Following protein blocking, specimens were incubated overnight at 4°C with the following primary antibodies: CD68 (1:100, MA5-16674, Invitrogen), Ki-67 (1:100, MA5-14520, Invitrogen), laminin (1:100, MA1-06100, Invitrogen), nMHC (1:100, PA5-72846, Invitrogen), sarcomeric α-actinin (1:100, ab9465, Abcam), Pax7 (1:100, PA1-117, Invitrogen), MyoD (1:100, MA1-41017, Invitrogen), CD206 (1:400, PA5-46994, Invitrogen) and CD80 (1:200, MA5-42562, Invitrogen). Following primary antibody incubation, specimens were washed with TBS-T (3 times, 5 min) and then incubated with the appropriate Alexa Fluor–conjugated secondary antibody (1:400, Invitrogen) for 2 hours at room temperature. Following secondary antibody incubation, specimens were washed with TBS-T (3 times, 5 min) and coverslips were mounted with Fluoroshield™ plus DAPI (Sigma-Aldrich) mounting medium. Immunofluorescent specimens were imaged using a multi-photon confocal fluorescence microscope (SP5, Leica) and quantitated using ImageJ software. Pooled data were generated by averaging data from 3-4 fields per specimen.

### Western Blot

Cells or tissues were washed with PBS and protein was extracted using RIPA Lysis and Extraction Buffer (Thermo Fisher) supplemented with Halt Protease and Phosphatase Inhibitor Cocktail (Thermo Fisher), according to the manufacturer’s recommended protocol. Protein concentration of the lysates was measured using the Pierce BCA protein assay kit (Thermo Fisher). Protein (20 µg) was separated, as previously described(67), by polyacrylamide gel electrophoresis using 4-12% Bis-Tris gels (Invitrogen) for 50 min at 200V and transferred to polyvinylidene difluoride membranes for 90 min at 250mA. Blots were then blocked with 5% non-fat dried milk (Bio-Rad) in TBS-T for 30 min at room temperature. Following protein blocking, blots were incubated overnight at 4°C with the following primary antibodies: TREX1 (1:1000, PA5-20665, Invitrogen & 1:400, #76892, Cell Signaling Technology; NPB1-76977, Novus Biologicals), Phospho-IKK alpha/beta (1:400, PA5-104696, Invitrogen), STING (1:500, ab288157, Abcam), Phospho-TBK1/NAK (1:400, #5483, Cell Signaling Technology), TBK1/NAK (1:400, #3504, Cell Signaling Technology), Phospho-STING (1:400, #72971, Cell Signaling Technology), cGAS (1:500, #31659, Cell signaling Technology), Pan-Actin (1:1000, #41185, Cell Signaling Technology), β-actin (1:1000, #12262, Cell Signaling Technology) and GAPDH (1:1000, #97166 & #51332, Cell Signaling Technology). Following primary antibody incubation, membranes were washed in TBS-T (3 times, 5 min) and then incubated in the appropriate HRP-conjugated secondary antibody (1:1000, #7074 & #7076, Cell Signaling Technology) for 2 hours at room temperature. Following secondary antibody incubation, blots were washed with TBS-T (3 times, 5 min) and imaged using Pierce ECL Western Blotting Substrate (Thermo Fisher) and a chemiluminescence ChemiDoc Imaging System (Bio-Rad). Protein expression was determined by densitometry using Image Lab (Bio-Rad).

### Flow Cytometry

Cells harvested from culture or mice were suspended in FACS buffer (2% FBS, 0.5M EDTA) at a concentration of 1 × 10^6^ cells per 100 µl. Cells were then incubated with FcR Blocking Reagent (Miltenyi Biotec, 130-092-575) for 10 min at 4°C and with incubation for 30 min on ice. After a PBS wash, cells were incubated 15 min on ice with Ghost Dye Red 710 (13-0871-T100, Tonbo Biosciences). Flow cytometry experiments were conducted using a SONY ID7000 Spectrum Cell Analyzer (Sony Biotechnology). Gates were established based on forward and side scatter to exclude cellular debris, and fluorescence compensation was carried out using single-labeled controls. Quantitative analysis was performed with ID7000 software (Sony Biotechnology, v2.0.2.17121).

### Macrophage Cytosolic Double-Stranded DNA Assay

Macrophages were grown on glass coverslips inside 6-well culture plates and processed for cytosolic double-stranded DNA (dsDNA) staining, as previously described(68). Briefly, macrophages were fixed with 4% paraformaldehyde for 10 min at room temperature, permeabilized with 0.1% saponin for 5 min at room temperature, and then incubated with protein blocking buffer for 1 hour at room temperature. Following protein blocking, macrophages were incubated overnight at 4°C with the following primary antibodies: dsDNA (1:1000, ab27156, Abcam) and COX-IV (1:400, ab16056, Abcam). Following primary antibody incubation, macrophages were washed with TBS-T (3 times, 5 min) and then incubated with the appropriate Alex Fluor-conjugated secondary antibody (1:400, Invitrogen) for 2 hours at room temperature. Following secondary antibody incubation, macrophages were washed with TBS-T (3 times, 5 min) and the cell-containing coverslips were mounted onto a Superfrost™ Plus microscope slides with Fluoroshield™ plus DAPI mounting medium. Immunofluorescent macrophages were imaged using a multi-photon confocal fluorescence microscope and quantitated using ImageJ software. Pooled data were generated by quantifying the number of dsDNA foci per cell and the percentage of cells containing dsDNA foci.

### Macrophage Phagocytosis Assay

To assess the phagocytic capacity of both unstimulated and stimulated macrophage populations, we conducted a fluorescent latex bead assay (Cayman Chemical), according to the manufacturer’s recommended protocol. Following M1 or M2 polarization, or RNA transfection, macrophages were exposed to FITC-labeled latex microspheres (0.1-μm diameter) for 2 hours in a cell incubator. Following the incubation, macrophages were detached and processed for flow cytometric analysis. Cell viability as measured with Ghost Dye Red 710 (13-0871-T100, Tonbo Biosciences) and quantification of microsphere uptake in each group by the percentage of FITC+ cells.

### Enzyme-linked Immunosorbent Assay

Protein lysates were collected from macrophages, conditioned media, or tissues, according to the manufacturer’s recommended protocol (Cayman Chemical) and processed for enzyme-linked immunosorbent assay (ELISA) to measure 2’3’-cGAMP levels (#501700, Cayman Chemical). Absorbance was measured at 450 nm with a microplate reader SpectraMax iD5 (Molecular devices).

### Myoblast Proliferation and Differentiation Assay

To measure the effect of 2’3’-cGAMP (MedChemExpress) on myoblast proliferation, C2C12 myoblasts were plated on 0.5% gelatin-coated plates in DMEM supplemented with 20% FBS and 1% penicillin/streptomycin and cultured until 30% confluency. C2C12 myoblasts were then transfected with 100 µM 2’3’-cGAMP using Lipofectamine RNAiMAX (Invitrogen), according to the manufacturer’s recommended protocol. Fresh 2’3’-cGAMP was added to the culture media daily for 3 consecutive days. After the third day, C2C12 myoblasts were processed for proliferation assay using the ClickTech Edu Cell Proliferation kit (Sigma-Aldrich), according to the manufacturer’s recommended protocol. Fluorescence was detected using the Cytation 5 cell imaging system (BioTek). EdU-positive and -negative nuclei were counted using Gen5 software (v3.08, Agilent). To measure the effect of 2’3’-cGAMP on differentiation, C2C12 myoblasts were cultured until 95% confluency and culture media was switched to DMEM supplemented with 2% FBS and 1% penicillin/streptomycin for 7 days. Fresh 2’3’-cGAMP (prepared as described above) was added to differentiation media each day for the first 3 consecutive days. On day 7, cells were fixed with 10% neutral buffered formalin for 15 min and permeabilized with 0.5% Triton X-100 (Sigma-Aldrich) for 20 min. Cells were washed with 3% bovine serum albumin (Sigma-Aldrich) and incubated with MYH4 antibody (1:400, #14-6503-82, Invitrogen) overnight at 4°C. Following primary antibody incubation, cells were washed with 3% bovine serum albumin and then incubated with Alexa Fluor 488 conjugated anti-mouse antibody (1:1000, Invitrogen) for 2 hours at room temperature. Following secondary antibody incubation, cells were washed with 3% bovine serum albumin and mounted using Fluoroshield + DAPI (Sigma-Aldrich). Fluorescence was detected using the Cytation 5 cell imaging system. Fusion was determined by the relative ratio of fused and unfused myonuclei.

### Statistical Analyses

Statistical significance was determined by independent *t*-tests (one-tailed), a one-way analysis of variance (ANOVA), or a two-way ANOVA, using GraphPad Prism v10. Normality was tested for all data sets, and the appropriate parametric test, as described above, was used. Significance was accepted at *P* ≤ 0.05. Data are represented as mean ± standard error of the mean (SEM).

## Data Availability

Resources may be made available upon reasonable request.

## AUTHOR CONTRIBUTIONS

Conceptualization: RGR, EM, and AI; Experimental Design: AR, RGR, EM; Data Collection: AR, NM, SC, AB, JZ, JA, LL, MF, WL, and SK; Data Analysis: AR, NM, JA; Manuscript Drafting: AR; Manuscript Editing: RGR, EM, and AI; Funding: RGR and EM; Final Approval: RGR.

## ACKNOWLEDGMENTS

The authors acknowledge the following Cedars-Sinai cores for their support: Applied Genomics, Computational and Translational Core; Biobank and Research Pathology Resource; and Flow Cytometry Core.

## FUNDING SUPPORT

Funding support was provided by NIH/NHLBI R01 HL167921 (to RGR) and NIH/NHLBI R01 HL164588 (to EM).

## CONFLICT OF INTERESTS

EM owns founder’s equity in Capricor. AI owns Capricor stock. Capricor has no relationship to this project, nor any licensing rights to the discoveries reported here. All other authors declare no competing interests.

## Notes

### Competing Interest Statement

EM owns founders equity in Capricor. AI owns Capricor stock. Capricor has no relationship to this project, nor any licensing rights to the discoveries reported here. All other authors declare no competing interests.

## REFERENCES

1. Romitti PA, Zhu Y, Puzhankara S, James KA, Nabukera SK, Zamba GK, et al. Prevalence of Duchenne and Becker muscular dystrophies in the United States. Pediatrics. 2015;135(3):513–21.

2. Schultz TI, Raucci Jr FJ, and Salloum FN. Cardiovascular disease in Duchenne muscular dystrophy: overview and insight into novel therapeutic targets. Basic to Translational Science. 2022;7(6):608–25.

3. Han R, Rader EP, Levy JR, Bansal D, and Campbell KP. Dystrophin deficiency exacerbates skeletal muscle pathology in dysferlin-null mice. Skeletal muscle. 2011;1:1–11.

4. Decout A, Katz JD, Venkatraman S, and Ablasser A. The cGAS–STING pathway as a therapeutic target in inflammatory diseases. Nature Reviews Immunology. 2021;21(9):548–69.

5. Ibrahim AG-E, Ciullo A, Komuro H, Miyamoto K, Jones XM, Yamaguchi S, et al. Augmentation of DNA exonuclease TREX1 in macrophages as a therapy for cardiac ischemic injury. Science Translational Medicine. 2025;17(827):eadp1338.

6. Miyamoto K, Jones XM, Yamaguchi S, Ciullo A, Li C, Coto JG, et al. Intravenous and oral administration of the synthetic RNA drug, TY1, reverses heart failure with preserved ejection fraction in mice. Basic Research in Cardiology. 2024:1–9.

7. Tulangekar A, and Sztal TE. Inflammation in duchenne muscular dystrophy– exploring the role of neutrophils in muscle damage and regeneration. Biomedicines. 2021;9(10):1366.

8. Yan N. Immune diseases associated with TREX1 and STING dysfunction. Journal of Interferon & Cytokine Research. 2017;37(5):198–206.

9. Paul BD, Snyder SH, and Bohr VA. Signaling by cGAS–STING in neurodegeneration, neuroinflammation, and aging. Trends in neurosciences. 2021;44(2):83–96.

10. Tripodi L, Villa C, Molinaro D, Torrente Y, and Farini A. The immune system in duchenne muscular dystrophy pathogenesis. Biomedicines. 2021;9(10):1447.

11. Pilling D, Galvis-Carvajal E, Karhadkar TR, Cox N, and Gomer RH. Monocyte differentiation and macrophage priming are regulated differentially by pentraxins and their ligands. BMC immunology. 2017;18:1–15.

12. Shim A, Luan X, Zhou W, Crow YJ, and Maciejowski J. Mutations in the non-catalytic polyproline motif destabilize TREX1 and amplify cGAS-STING signaling. Human Molecular Genetics. 2024:ddae089.

13. McDonald CM, Henricson EK, Abresch RT, Duong T, Joyce NC, Hu F, et al. Long-term effects of glucocorticoids on function, quality of life, and survival in patients with Duchenne muscular dystrophy: a prospective cohort study. The Lancet. 2018;391(10119):451–61.

14. Wang N, Wang X, Li Y, Shen H, Liu Z, Ma Z, et al. The STING-IRF3 pathway contributes to paraquat-induced acute lung injury. Toxicology Mechanisms and Methods. 2022;32(2):145–57.

15. Quek H, Luff J, Cheung K, Kozlov S, Gatei M, Lee CS, et al. A rat model of ataxia-telangiectasia: evidence for a neurodegenerative phenotype. Human molecular genetics. 2017;26(1):109–23.

16. Vico T, Youssif C, Zare F, Comalada M, Sebastian C, Lloberas J, and Celada A. GM-CSF protects macrophages from DNA damage by inducing differentiation. Cells. 2022;11(6):935.

17. Colonna M. DNA damage response impacts macrophage functions. *Blood*, The Journal of the American Society of Hematology. 2015;126(22):2440–2.

18. Füchtbauer AF, Preus S, Börjesson K, McPhee SA, Lilley DM, and Wilhelmsson LM. Fluorescent RNA cytosine analogue–an internal probe for detailed structure and dynamics investigations. Scientific reports. 2017;7(1):2393.

19. Pereira-Lopes S, Celhar T, Sans-Fons G, Serra M, Fairhurst A-M, Lloberas J, and Celada A. The exonuclease Trex1 restrains macrophage proinflammatory activation. The Journal of Immunology. 2013;191(12):6128–35.

20. Cao DJ, Schiattarella GG, Villalobos E, Jiang N, May HI, Li T, et al. Cytosolic DNA sensing promotes macrophage transformation and governs myocardial ischemic injury. Circulation. 2018;137(24):2613–34.

21. Zhang Y, Zhang C, Feng R, Meng T, Peng W, Song J, et al. CXCR4 regulates macrophage M1 polarization by altering glycolysis to promote prostate fibrosis. Cell Communication and Signaling. 2024;22(1):456.

22. Guillemot-Legris O, and Muccioli GG. The oxysterome and its receptors as pharmacological targets in inflammatory diseases. British journal of pharmacology. 2022;179(21):4917–40.

23. Young LV, Wakelin G, Cameron AW, Springer SA, Ross JP, Wolters G, et al. Muscle injury induces a transient senescence-like state that is required for myofiber growth during muscle regeneration. The FASEB Journal. 2022;36(11):e22587.

24. Riuzzi F, Beccafico S, Sagheddu R, Chiappalupi S, Giambanco I, Bereshchenko O, et al. Levels of S100B protein drive the reparative process in acute muscle injury and muscular dystrophy. Scientific reports. 2017;7(1):12537.

25. de Masson A, Darbord D, Dobos G, Boisson M, Roelens M, Ram-Wolff C, et al. Macrophage-derived CXCL9 and CXCL11, T-cell skin homing, and disease control in mogamulizumab-treated CTCL patients. *Blood*, The Journal of the American Society of Hematology. 2022;139(12):1820–32.

26. Kim J-Y, Cho D-W, Choi J-Y, Jeong S, Kang M, Kim WJ, et al. CXCL11 reprograms M2-biased macrophage polarization to alleviate pulmonary fibrosis in mice. Cell & Bioscience. 2024;14(1):140.

27. Altemeier WA, Zhu X, Berrington WR, Harlan JM, and Liles WC. Fas (CD95) induces macrophage proinflammatory chemokine production via a MyD88-dependent, caspase-independent pathway. Journal of Leucocyte Biology. 2007;82(3):721–8.

28. Kobayashi T, Matsuoka K, Sheikh SZ, Elloumi HZ, Kamada N, Hisamatsu T, et al. NFIL3 is a regulator of IL-12 p40 in macrophages and mucosal immunity. The Journal of Immunology. 2011;186(8):4649–55.

29. Okamoto T, Gohil K, Finkelstein EI, Bove P, Akaike T, and Van Der Vliet A. Multiple contributing roles for NOS2 in LPS-induced acute airway inflammation in mice. American Journal of Physiology-Lung Cellular and Molecular Physiology. 2004;286(1):L198–L209.

30. Burgess M, Wicks K, Gardasevic M, and Mace KA. Cx3CR1 expression identifies distinct macrophage populations that contribute differentially to inflammation and repair. Immunohorizons. 2019;3(7):262–73.

31. Liang F, Giordano C, Shang D, Li Q, and Petrof BJ. The dual CCR2/CCR5 chemokine receptor antagonist Cenicriviroc reduces macrophage infiltration and disease severity in Duchenne muscular dystrophy (Dmdmdx-4Cv) mice. PLoS One. 2018;13(3):e0194421.

32. Patsalos A, Halasz L, Medina-Serpas MA, Berger WK, Daniel B, Tzerpos P, et al. A growth factor–expressing macrophage subpopulation orchestrates regenerative inflammation via GDF-15. Journal of Experimental Medicine. 2021;219(1):e20210420.

33. Yadav P, Ortega JG, Tamaki W, Chien C, Chang K-c, Biswas N, et al. Macrophage-fibroblast crosstalk drives Arg1-dependent lung fibrosis via ornithine loading. bioRxiv. 2023:2023.09. 06.556606.

34. Di Carmine S, Scott MM, McLean MH, and McSorley HJ. The role of interleukin-33 in organ fibrosis. Discovery Immunology. 2022;1(1):kyac006.

35. Jiao B, Zhang Q, Jin C, Yu H, and Wu Q. IRF4 participates in pulmonary fibrosis induced by silica particles through regulating macrophage polarization and fibroblast activation. Inflammation. 2024;47(1):45–59.

36. Knight DA, Ernst M, Anderson GP, Moodley YP, and Mutsaers SE. The role of gp130/IL-6 cytokines in the development of pulmonary fibrosis: critical determinants of disease susceptibility and progression? Pharmacology & therapeutics. 2003;99(3):327–38.

37. Rogers RG, Fournier M, Sanchez L, Ibrahim AG, Aminzadeh MA, Lewis MI, and Marbán E. Disease-modifying bioactivity of intravenous cardiosphere-derived cells and exosomes in mdx mice. JCI insight. 2019;4(7).

38. McDonald CM, Marbán E, Hendrix S, Hogan N, Smith RR, Eagle M, et al. Repeated intravenous cardiosphere-derived cell therapy in late-stage Duchenne muscular dystrophy (HOPE-2): a multicentre, randomised, double-blind, placebo-controlled, phase 2 trial. The Lancet. 2022;399(10329):1049–58.

39. Lynch GS, Hinkle RT, Chamberlain JS, Brooks SV, and Faulkner JA. Force and power output of fast and slow skeletal muscles from mdx mice 6-28 months old. The Journal of physiology. 2001;535(2):591–600.

40. Van Erp C, Loch D, Laws N, Trebbin A, and Hoey AJ. Timeline of cardiac dystrophy in 3–18-month-old MDX mice. Muscle & nerve. 2010;42(4):504–13.

41. Aminzadeh MA, Rogers RG, Fournier M, Tobin RE, Guan X, Childers MK, et al. Exosome-mediated benefits of cell therapy in mouse and human models of Duchenne muscular dystrophy. Stem cell reports. 2018;10(3):942–55.

42. Schmidt WM, Uddin MH, Dysek S, Moser-Thier K, Pirker C, Höger H, et al. DNA damage, somatic aneuploidy, and malignant sarcoma susceptibility in muscular dystrophies. PLoS genetics. 2011;7(4):e1002042.

43. Tidball JG. Mechanisms of muscle injury, repair, and regeneration. Comprehensive physiology. 2011(4):2029–62.

44. Sun J, Li L, Hu J, Gao Y, Song J, Zhang X, and Hu H. Time-course RNA-Seq profiling reveals isoform-level gene expression dynamics of the cGAS-STING pathway. Computational and Structural Biotechnology Journal. 2022;20:6490–500.

45. Shams AS, Arpke RW, Gearhart MD, Weiblen J, Mai B, Oyler D, et al. The chemokine receptor CXCR4 regulates satellite cell activation, early expansion, and self-renewal, in response to skeletal muscle injury. Frontiers in Cell and Developmental Biology. 2022;10:949532.

46. Zhou S, Li T, Zhang W, Wu J, Hong H, Quan W, et al. The cGAS-STING-interferon regulatory factor 7 pathway regulates neuroinflammation in Parkinson’s disease. Neural Regeneration Research. 2025;20(8):2361–72.

47. Elias M, Zhao S, Le HT, Wang J, Neurath MF, Neufert C, et al. IL-36 in chronic inflammation and fibrosis—bridging the gap? The Journal of clinical investigation. 2021;131(2).

48. Onódi Z, Szabó PL, Kucsera D, Pokreisz P, Dostal C, Hilber K, et al. Inflammasome activity in the skeletal muscle and heart of rodent models for Duchenne muscular dystrophy. International Journal of Molecular Sciences. 2023;24(10):8497.

49. Mi S, Huang F, Jiao M, Qian Z, Han M, Miao Z, and Zhan H. Inhibition of MEG3 ameliorates cardiomyocyte apoptosis and autophagy by regulating the expression of miRNA-129-5p in a mouse model of heart failure. Redox Report. 2023;28(1):2224607.

50. Li J, Song Y, Jin J-Y, Li G-H, Guo Y-Z, Yi H-Y, et al. CD226 deletion improves post-infarction healing via modulating macrophage polarization in mice. Theranostics. 2020;10(5):2422.

51. Yutzey KE. Neuregulin 1 makes heart muscle. Nature. 2015;520(7548):445–6.

52. Cheng YY, Yan YT, Lundy DJ, Lo AH, Wang YP, Ruan SC, et al. Reprogramming-derived gene cocktail increases cardiomyocyte proliferation for heart regeneration. EMBO molecular medicine. 2017;9(2):251–64.

53. Petrof BJ. Macrophage plasticity in Duchenne muscular dystrophy: a nexus of pathological remodelling with therapeutic implications. The Journal of Physiology. 2022;600(15):3455–64.

54. Burnett SH, Kershen EJ, Zhang J, Zeng L, Straley SC, Kaplan AM, and Cohen DA. Conditional macrophage ablation in transgenic mice expressing a Fas-based suicide gene. Journal of Leucocyte Biology. 2004;75(4):612–23.

55. Zegard A, Okafor O, De Bono J, Kalla M, Lencioni M, Marshall H, et al. Myocardial fibrosis as a predictor of sudden death in patients with coronary artery disease. Journal of the American College of Cardiology. 2021;77(1):29–41.

56. Summan M, Warren GL, Mercer RR, Chapman R, Hulderman T, Van Rooijen N, and Simeonova PP. Macrophages and skeletal muscle regeneration: a clodronate-containing liposome depletion study. *American Journal of Physiology-Regulatory*, Integrative and Comparative Physiology. 2006;290(6):R1488–R95.

57. Zhou W, Richmond-Buccola D, Wang Q, and Kranzusch PJ. Structural basis of human TREX1 DNA degradation and autoimmune disease. Nature communications. 2022;13(1):4277.

58. Li T, and Chen ZJ. The cGAS–cGAMP–STING pathway connects DNA damage to inflammation, senescence, and cancer. Journal of Experimental Medicine. 2018;215(5):1287–99.

59. Bostick B, Yue Y, and Duan D. Gender influences cardiac function in the mdx model of Duchenne cardiomyopathy. Muscle & nerve. 2010;42(4):600–3.

60. Ibrahim AG-E, Ciullo A, Miyamoto K, Liao K, Jones X, Yamaguchi S, et al. Augmentation of DNA exonuclease TREX1 in macrophages as a therapy for cardiac ischemic injury. Biorxiv. 2024:2024.02. 20.581294.

61. Rogers RG, Baumann CW, and Otis JS. Recovery of skeletal muscle function following injury is not augmented by acute resveratrol supplementation. Int J Clin Exp Physiol. 2015;2(1):29.

62. Rogers RG, Li L, Peck K, Sanchez L, Liu W, Ciullo A, et al. Cardiosphere-derived cells, with and without a biological scaffold, stimulate myogenesis and recovery of muscle function in mice with volumetric muscle loss. Biomaterials. 2021;274:120852.

63. Hong Y, Rannou A, Manriquez N, Antich J, Liu W, Fournier M, et al. Cardiac and skeletal muscle manifestations in the G608G mouse model of Hutchinson-Gilford progeria syndrome. Aging Cell. 2024;23(10):e14259.

64. Baumann CW, Rogers RG, and Otis JS. Utility of 17-(allylamino)-17-demethoxygeldanamycin treatment for skeletal muscle injury. Cell Stress and Chaperones. 2016;21(6):1111–7.

65. De Couto G, Jaghatspanyan E, DeBerge M, Liu W, Luther K, Wang Y, et al. Mechanism of enhanced MerTK-dependent macrophage efferocytosis by extracellular vesicles. Arteriosclerosis, thrombosis, and vascular biology. 2019;39(10):2082–96.

66. Rogers RG, and Otis JS. Resveratrol-mediated expression of KLF15 in the ischemic myocardium is associated with an improved cardiac phenotype. Cardiovascular drugs and therapy. 2017;31:29–38.

67. Sanchez L, Mesquita T, Zhang R, Liao K, Rogers R, Lin Y-N, et al. MicroRNA-dependent suppression of biological pacemaker activity induced by TBX18. Cell Reports Medicine. 2022;3(12).

68. Sato A, Buque A, Yamazaki T, Bloy N, Petroni G, and Galluzzi L. Immunofluorescence microscopy-based assessment of cytosolic DNA accumulation in mammalian cells. STAR protocols. 2021;2(2):100488.

