## Supplemental Data for "Pathogenic role of cGAS-STING pathway in Duchenne muscular dystrophy"

Alice Rannou *et al.*

Corresponding author: Russell G. Rogers, PhD

**The PDF file includes:**

Figs. S1 to S6
Table S1

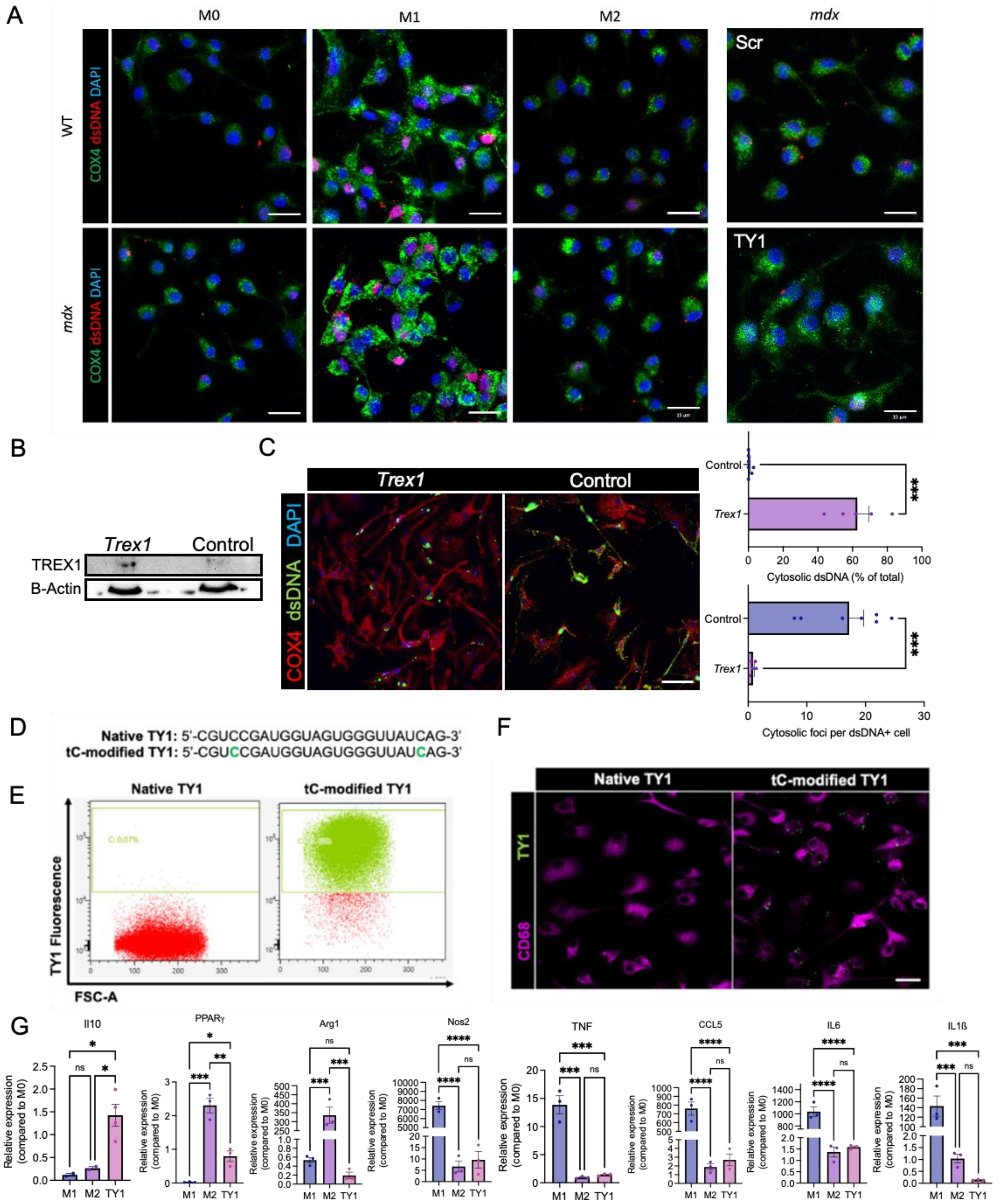


**Supplemental Figure 1: Characteristics of mouse macrophages.** Wild-type and *mdx* bone marrow derived-macrophages were isolated and cultured for 7 days in presence of m-CSF (10 ng/mL). At day 7, M0 (inactive) macrophages were polarized to M1 (LPS 0.1 µg/ml; IFN-γ 50 ng/mL), M2 (IL13 10 ng/mL; IL4 10 ng/mL), or conditioned by TY1 or scramble (50 nM) for 24 hours. The percentage of cells containing cytosolic dsDNA+ foci and the number of positive dsDNA+ foci per cell were quantified (**A**). *Mdx* bone marrow derived-macrophages were isolated and cultured for 7 days in presence of m-CSF (10 ng/mL). At day 7, macrophages (M0) were polarized to M1 (LPS 0.1 μg/ml; IFN-γ 50 ng/mL) and then transfected with *Trex1* plasmid DNA or control. Western blot shows *Trex1* plasmid DNA increased TREX1 protein levels (**B**). Immunocytochemistry shows *Trex1* plasmid DNA decreased accumulation of cytosolic double-stranded DNA (dsDNA) (**C**). *mdx* bone marrow derived-macrophages were isolated and cultured for 7 days in presence of m-CSF (10 ng/mL). At day 7, macrophages (M0) were conditioned by a fluorescent TY1 (50 nM) for 24 hours to confirm macrophage uptake. The modifications of the native sequence of TY1 are in green and indicate the tC-modified bases (**D**). After 24h, cells were collected, and fluorescence was measured by flow cytometry (**E**) and by confocal microscopy (**F**). qPCR of IL10, TGFß1, Nos2, PPARγ, TNF, CCL5, IL6, IL1ß and Arg1 in M1 (pro-inflammatory), M2 (anti-inflammatory) and macrophages conditioned by TY1 (**E**). Scale bar: 15 μm. N = 2-3 per group. Data are represented as mean ± SEM. Statistical significance was determined by ANOVA. *$p < 0.05$, **$p < 0.01$, ***$p < 0.001$, ****$p < 0.0001$, ns = not significant.

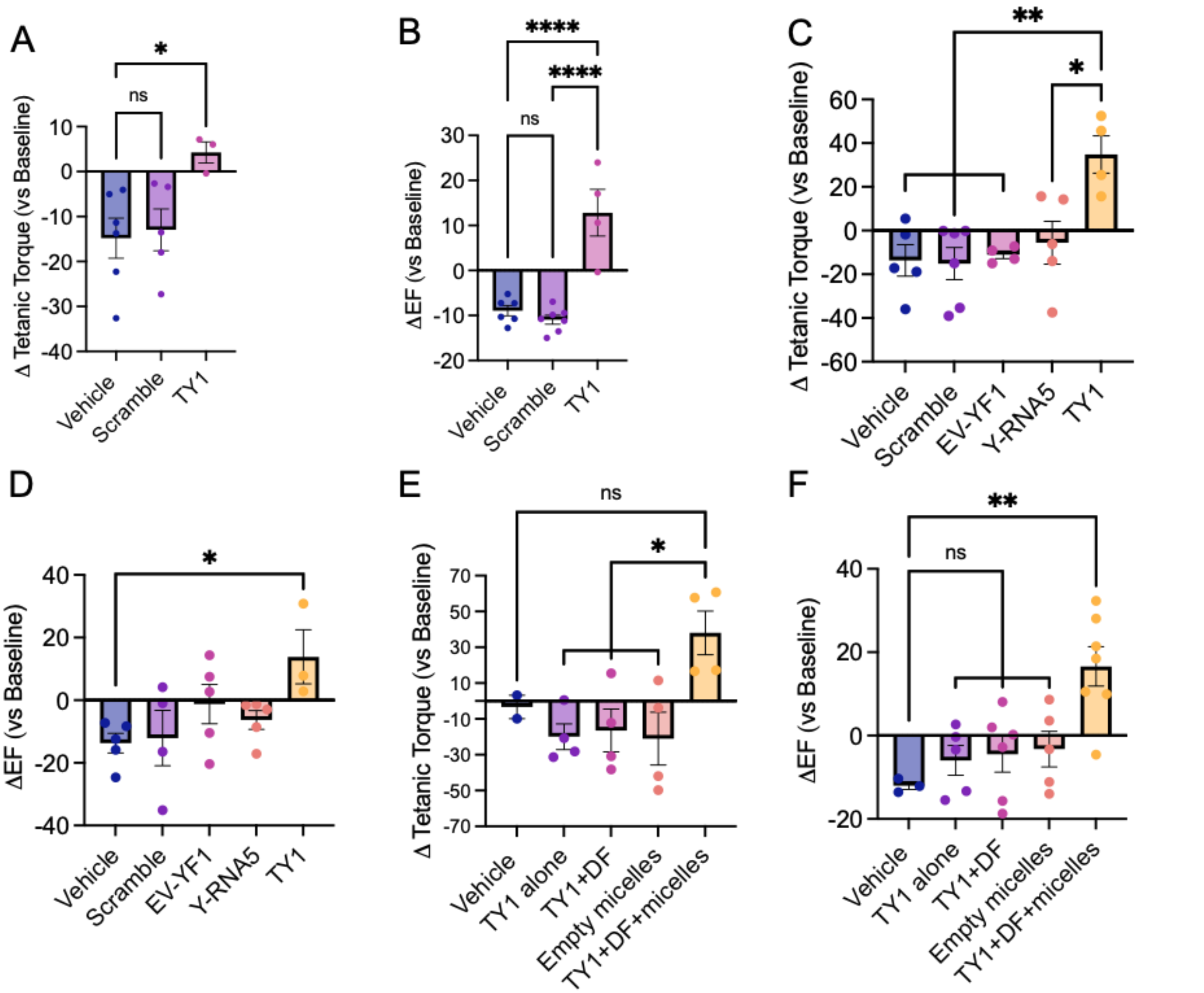


**Supplemental Figure 2: Benefits of TY1 are unique to its sequence and formulation.** *Mdx* mice received vehicle, scramble, or TY1 (**A&B**); vehicle, scramble EV-YF1, Y-RNA5, or TY1 (**C&D**); or vehicle, TY1 alone, TY1+DharmaFECT (DF), empty micelles, or TY1+DF+micelles (**E&F**), twice-weekly via gavage administration for 8 weeks. All RNAs were formulated at 0.2 μg/kg body weight. Functional measurements were made at baseline and endpoint. Isometric tetanic torque was only improved by TY1 when formulated with DF and micelles (**A, C, E**). Likewise, ejection fraction was only improved by TY1 when formulated with DF and micelles (**B, D, F**). N = 3-7 per group. Data are represented as mean ± SEM. Statistical significance was determined by ANOVA. *p < 0.05, **p < 0.01, ***p < 0.001, ****p < 0.0001, ns = not significant.

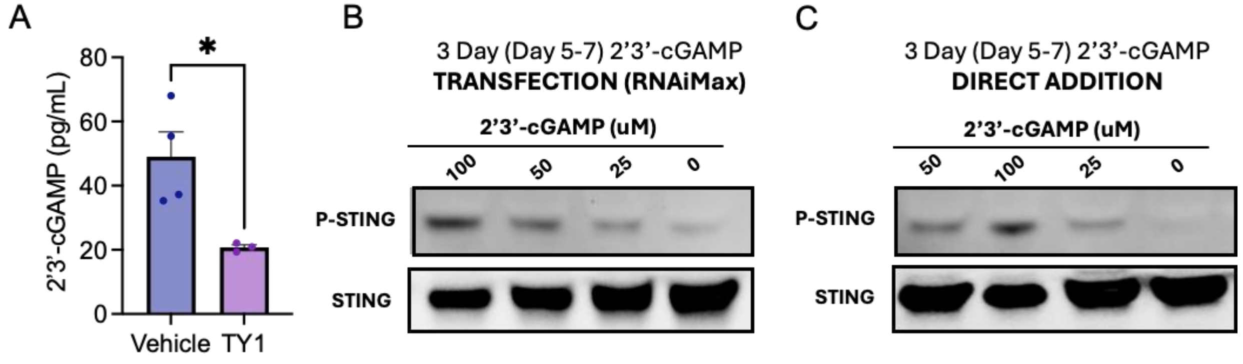


**Supplemental Figure 3: 2'3'-cGAMP secretion and STING phosphorylation.** *Mdx* bone marrow derived-macrophages were isolated and cultured for 7 days in presence of m-CSF (10 ng/mL). At day 7, macrophages (M0) were conditioned by TY1 (50 nM) for 24 hours. ELISA shows TY1 lowers the secretion of 2'3'-cGAMP into the cell culture media (**A**). C2C12 myoblasts were cultured and exposed to varying concentrations of 2'3'-cGAMP. Western blot shows 2'3'-cGAMP increases STING phosphorylation as a function of concentration whether packaged into a transfection reagent (**B**) or not (**C**). N = 3-4 per group. Data are represented as mean ± SEM. Statistical significance was determined by independent *t*-test. *p < 0.05.

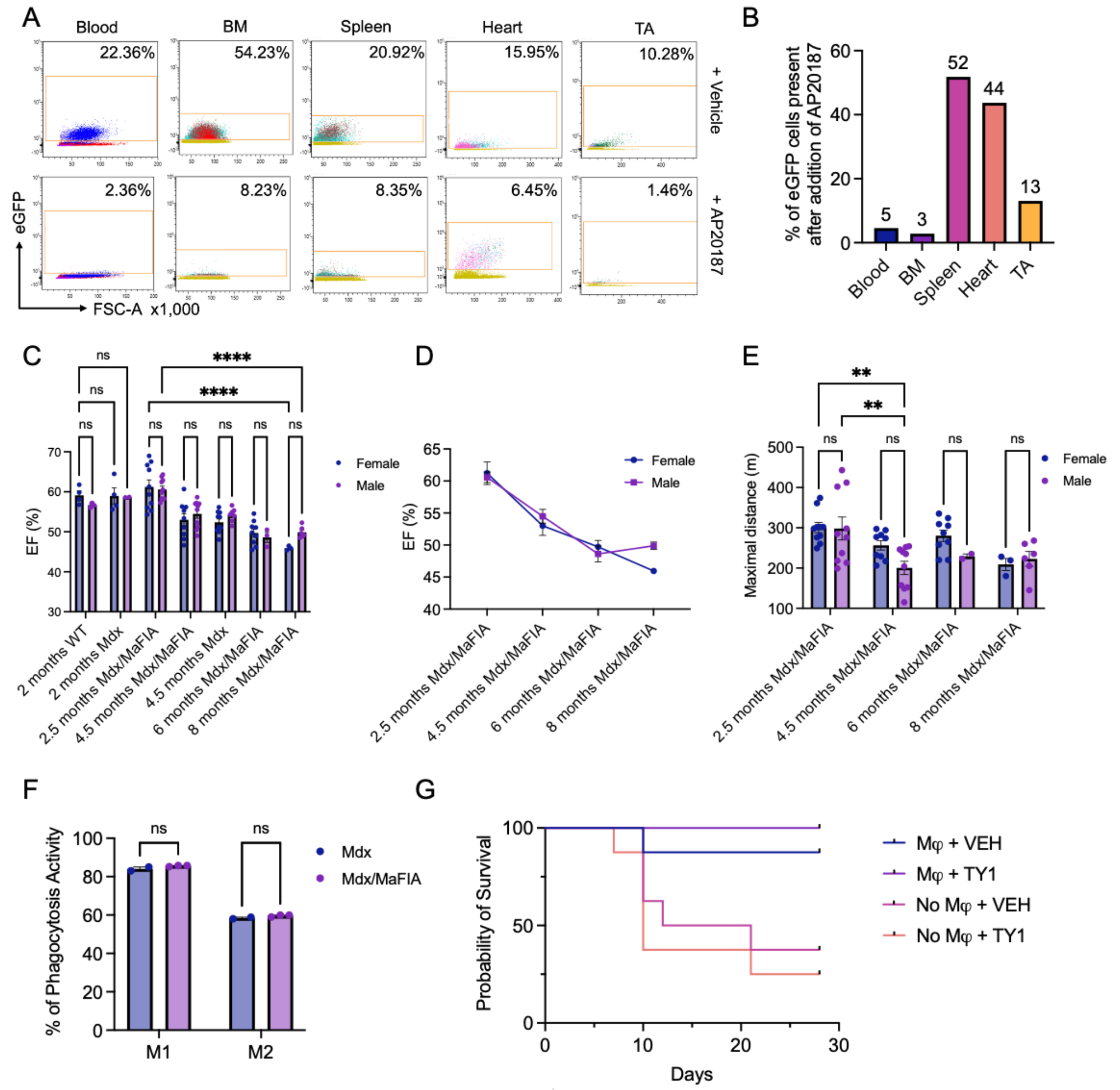


**Supplemental Figure 4: Characterization of macrophage-depletable *mdx* mice.** A macrophage-depletable *mdx* mouse was generated by crossing *mdx* and MaFIA mice (*mdx*/MaFIA). When administered AP20187, monocytes in the blood and bone marrow and macrophages in spleen, heart, and TA were depleted, as measured by flow cytometry (**A&B**). Macrophages from *mdx*/MaFIA mice express eGFP, enabling fluorescent tracking. Ejection fraction was not different from wild-type or *mdx* mice at 2 months of age but declined with age, consistent with progression of cardiomyopathy in *mdx* mice (**C&D**). No notable differences were observed between males and females in heart function (**D**) or exercise capacity (**E**). The phagocytic capacity of M1 and M2 macrophages were not different between *mdx* and *mdx*/MaFIA mice (**F**). Probability of survival of *mdx*/MaFIA mice with or without AP20187 (**G**). N = 2-10 per group. Data are represented as mean ±

SEM. Statistical significance was determined by ANOVA. $*p < 0.05$, $**p < 0.01$, $***p < 0.001$, $****p < 0.0001$, ns = not significant.

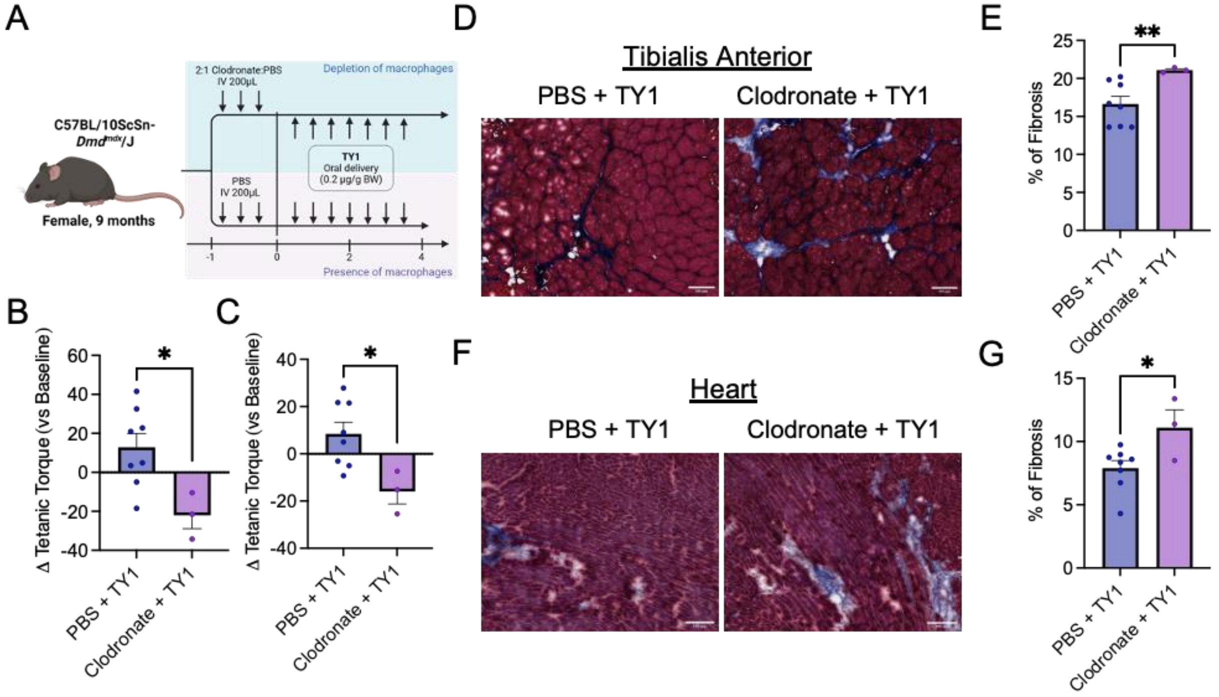


**Supplemental Figure 5: Characterization of macrophage depletion in *mdx* mice.** Schematic of experimental design (**A**). *Mdx* mice were given three consecutive loading injections (200μL, intravenous) of clodronate or PBS. TY1 (0.2 μg/g body weight) was given via oral gavage twice a week until endpoint. Functional measurements were made at baseline and endpoint. Heart and tibialis anterior (TA) were isolated at endpoint for histological assessment. Clodronate administration eliminated TY1's improvement in isometric tetanic torque (**B**) and ejection fraction (**C**). Masson Trichrome analysis revealed clodronate eliminated TY1's anti-fibrotic properties in both TA (**D&E**) and Heart (**F&G**) tissue. Scale Bars = 100 μM. N = 3-8 per group. (Clodronate + TY1), N = 8 (PBS + TY1). Data are represented as mean ± SEM. Statistical significance was determined by independent *t*-test. *P<0.05, **P<0.01.

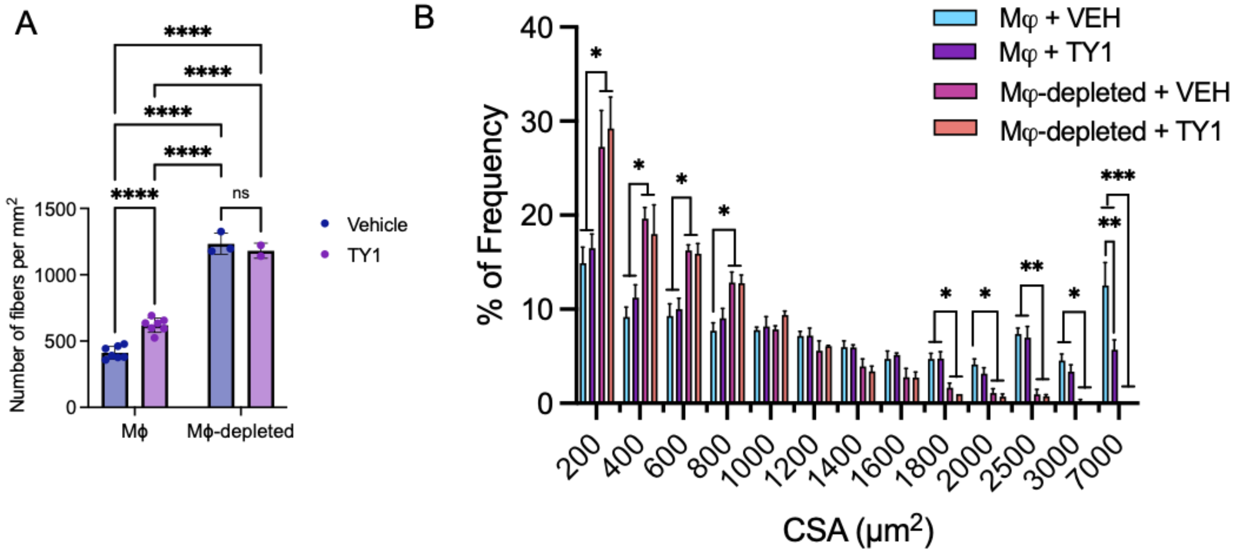


**Supplemental Figure 6: Effects of macrophage depletion on myofiber size.** Analysis of histological specimens from *mdx*/MaFIA mouse tibialis anterior (TA) demonstrate the effects of macrophages and TY1 on myofiber numbers and size. TY1 boosts the number of myofibers but only when macrophages are present (**A**). Depleting macrophages, regardless of intervention, led to greater number of myofibers but the myofibers were smaller than those from macrophage-competent mice (**B**). No major differences in myofiber size frequency distribution by TY1 (**B**). N = 2-6 per group. Bar graphs: mean ± SEM. Statistical significance was determined by ANOVA. *p < 0.05, **p < 0.01, ***p < 0.001, ****p < 0.0001, ns = not significant.

| Gene names | Primer sequences | |
|---|---|---|
| CCL5 | F<br>R | ATATGGCTCGGACACCACTC<br>ACTTGGCGGTTCCTTCGAG |
| IL1ß | F<br>R | TGCCACCTTTTGACAGTGATG<br>AAGGTCCACGGGAAAGACAC |
| IL1α | F<br>R | ACGTCAAGCAACGGGAAGAT<br>AAGGTGCTGATCTGGGTTGG |
| Nos2 | F<br>R | CTATGGCCGCTTTGATGTGC<br>CAACCTTGGTGTTGAAGGCG |
| TGFß1 | F<br>R | CATCCATGACATGAACCGGC<br>GAAGTTGGCATGGTAGCCCT |
| PPARγ | F<br>R | AAGAGCTGACCCAATGGTTG<br>GGATCCGGCAGTTAAGATCA |
| Arg1 | F<br>R | AGCACTGAGGAAAGCTGGTC<br>TACGTCTCGCAAGCCAATGT |
| IL6 | F<br>R | GGGACTGATGCTGGTGACAA<br>ACAGGTCTGTTGGGAGTGGT |
| TNF | F<br>R | TAGCCCACGTCGTAGCAAAC<br>ACAAGGTACAACCCATCGGC |
| IL10 | F<br>R | GGCCCAGAAATCAAGGAGCA<br>CACACTGCAGGTGTTTTAGCTT |

**Supplemental Table 1:**

Sequences of the forward (F) and reverse (R) primers used for qPCR for CCL5, IL1ß, IL1α, Nos2, TGFß1, PPARγ, Arg1, IL6, TNF and IL10.
